# HDAC1/2 maintain *TERT* mRNA stability in *TERT*-rearranged neuroblastoma

**DOI:** 10.64898/2026.11.15.766789

**Authors:** Sabine Finkler, Annabell Szymansky, Annika Sprüssel, Sveva Castelli, Charlotte Gatzweiler, Franziska Schulze, Jan F. Hollander, Rocío Chamorro González, Kathrin Hauptmann, Steffen Fuchs, Konstantin Helmsauer, Angelika Eggert, Dieter Beule, Anja Kühl, Marco Lodrini, Anton G. Henssen, Nils Blüthgen, January Weiner, Hedwig E. Deubzer

## Abstract

Genomic *TERT* locus rearrangements define an aggressive high-risk neuroblastoma subgroup characterized by enhancer hijacking, high-level *TERT* expression and continuous telomerase activation. Although these structural alterations supply a strong transcriptional drive, the post-transcriptional mechanisms preserving *TERT* mRNA presence remain incompletely understood. We identified histone deacetylases 1 and 2 (HDAC1/2) as essential post-transcriptional guardians of *TERT* mRNA stability in *TERT*-rearranged neuroblastoma. Pharmacological inhibition or RNAi-mediated depletion of HDAC1 and HDAC2, but not HDACs 3-11, rapidly downregulated *TERT* transcript levels and suppressed telomerase enzymatic activity. Transcript decay kinetics following transcriptional arrest with actinomycin D confirmed that HDAC1/2 blockade accelerates *TERT* mRNA decay, reducing transcript half-life by 30-47%. High-resolution ChIP-seq profiling across the translocation breakpoint demonstrated HDAC inhibition does not alter local chromatin accessibility. Array-based DNA methylation profiling showed only a mild, non-specific global reduction in CpG island methylation upon panobinostat exposure and no focal correlation with *TERT* transcript loss across the breakpoint locus. Depleting *TERT* mRNA via HDAC1/2 inhibition dismantled a core 50-gene *TERT*-associated oncogenic expression network. Ectopic re-expression of *TERT* cDNA partially rescued cellular viability and normalized expression of key signature genes despite continuous HDAC inhibition. HDAC1/2 inhibition significantly attenuated tumor growth, reduced intratumor *TERT* expression and prolonging overall survival in both preventive and therapeutic *TERT*-rearranged xenograft mouse models. Our data establish a post-transcriptional rather than transcriptional mode of *TERT* regulation and confirm tumor suppression is partly driven by depleting *TERT* transcripts, uncovering a HDAC1/2 dependency in *TERT*-driven neuroblastoma. We provide a molecular rationale for therapeutic strategies disrupting HDAC1/2-mediated *TERT* mRNA stabilization.

## INTRODUCTION

Neuroblastoma is a pediatric solid tumor of the sympathetic nervous system derived from neural crest progenitor cells, representing the most common extracranial malignancy in infancy and early childhood. Accounting for approximately 13 cases per million children annually in industrialized nations and nearly 15% of all pediatric cancer-related deaths worldwide, neuroblastoma exhibits extreme clinical and biological diversity.^1^ Clinical behavior ranges from complete spontaneous regression in low-risk infant tumors^2^ to widespread metastasis and chemotherapy-resistant fatal relapse in high-risk disease^3^.

Clinical management relies on rigorous risk stratification^4^ incorporating patient age, International Neuroblastoma Risk Group (INRG)^5^ stage, histological classification and key driver genomic alterations^6^. Despite aggressive, multimodal therapeutic protocols – combining intensive induction polychemotherapy, surgical resection, high-dose consolidation chemotherapy with autologous stem cell rescue, radiotherapy and targeted anti-GD2 immunotherapy - the prognosis for patients with high-risk neuroblastoma remains unsatisfactory, with 5-year overall survival stagnating around 50%.^7^ Approximately 10% of high-risk patients suffer from primary refractory disease, and relapsed cases carry acquired resistance to conventional cytotoxic regimens.^8^ Long-term survivors endure profound, lifelong treatment-related organ toxicities^1^, emphasizing the imperative to identify precise molecular targets and develop mechanistically driven therapeutic interventions.

A cornerstone of neuroblastoma oncogenesis is the acquisition of telomere maintenance mechanisms, which confer cellular immortality and drive aggressive clinical phenotypes.^9^ Whole-genome sequencing and molecular profiling studies established that high-risk neuroblastomas almost universally harbor one of three mutually exclusive, non-overlapping telomere maintenance mechanisms: *MYCN* amplification (∼40-50% of high-risk cases)^10,11^, genomic rearrangements involving the telomerase reverse transcriptase (*TERT*) locus (∼20-30%)^12,13^, or loss-of-function aberrations in the chromatin remodeler *ATRX* (∼10-15%)^14,15^, which trigger the alternative lengthening of telomeres (ALT) pathway.

Genomic rearrangements at the *TERT* locus (chromosome 5p15.33) define a distinct, highly aggressive subgroup of high-risk neuroblastomas characterized by elevated *TERT* transcript levels, high enzymatic telomerase activity and dismal patient outcomes.^12,13^ Unlike adult carcinomas and glioblastomas, in which somatic hotspot single-nucleotide point mutations in the *TERT* promoter predominate (e.g., *C228T*, *C250T*), *TERT* promoter mutations are exceedingly rare in neuroblastoma.^16,17^ Instead, *TERT* activation in pediatric neuroblastoma is driven almost exclusively by structural genomic variants including chromosomal translocations, inversions and intrachromosomal deletions.^12,13^ These structural rearrangements alter the local 3D chromatin architecture by juxtaposing the coding sequence of *TERT* to strong, distal super-enhancers, a mechanism known as ‘enhancer hijacking’.^12,18^ Enhancer hijacking leads to massive local chromatin remodeling, high-level *TERT* transcription, and productive transcriptional elongation driven by bromodomain protein Brd4 and cyclin-dependent kinases.^19^ Functional transcriptomic profiling further revealed that high *TERT* expression in non-*MYCN*-amplified neuroblastoma is accompanied by the coordinated activation of a core 50-gene TERT-associated oncogenic expression network.^19^ This signature includes critical cell cycle drivers, DNA repair components and mitotic regulators such as AURKA, BIRC5 (survivin), E2F8, FEN1, EXO1, MIS18A, UHRF1 and CDCA5, which together sustain tumor cell survival and therapeutic resistance.^19^

Because the catalytic subunit of telomerase is active in 80-90 % of human malignancies but repressed in most adult somatic tissues, TERT has long represented a high-priority therapeutic target across oncology.^20,21^ Over more than two decades, intense drug development efforts have explored multiple strategies to block telomerase, including direct catalytic nucleoside analogues, G-quadruplex stabilizers, oligonucleotide competitive inhibitors (such as imetelstat/GRN163L), and TERT-targeted immunotherapies.^22^ Despite robust preclinical antitumoral activity, direct telomerase inhibitors have systematically failed to achieve clinical translation due to major pharmacological bottlenecks and systemic toxicities.^22,23^ Direct catalytic inhibition requires multi-generational cell divisions to induce critical telomere shortening before triggering senescence or apoptosis, during which tumor cells frequently acquire resistance mechanisms.^22^ Furthermore, clinical evaluation of agents like imetelstat in pediatric solid and CNS tumors revealed severe, dose-limiting hematologic toxicities, including grade 3/4 thrombocytopenia, lymphopenia, and neutropenia.^23^ Given that no direct TERT inhibitors are clinically available, identifying indirect, actionable druggable dependencies that rapidly suppress *TERT* expression and dismantle telomerase activity represents a crucial unmet clinical need in (pediatric) oncology. We here demonstrate that class I histone deacetylases 1 and 2 (HDAC1/2) act as essential post-transcriptional guardians that actively protect *TERT* mRNA from rapid degradation, providing a compelling molecular basis for indirect therapeutic intervention.

## RESULTS

### HDAC1 and HDAC2 regulate *TERT* transcript levels, telomerase activity and transcript stability in *TERT*-rearranged neuroblastoma cells

To investigate whether histone deacetylase (HDAC) activity is required to maintain elevated *TERT* transcript levels in neuroblastoma cells harboring genomic rearrangements involving the *TERT* locus (5p15.33) followed by enhancer hijacking, high-level *TERT* expression and continuous telomerase activation, we treated GI-ME-N and CLB-GA cells with low nanomolar concentrations of the FDA- and EMA-approved pan-HDAC inhibitor panobinostat. Because GI-ME-N and CLB-GA represent two cell culture models^12^ known to harbor endogenous *TERT* rearrangements, they served as the preclinical models for this study. Targeted hybrid capture panel sequencing^24^ across key pediatric neuroblastoma driver genes confirmed their baseline genomic profiles (**Suppl. Fig. S1**). Quantitative RT-PCR analysis demonstrated a rapid, time-dependent reduction in *TERT* mRNA levels, after 2 h of treatment in GI-ME-N cells and after 4 h in CLB-GA cells (**Fig. 1a**). At maximal suppression, *TERT* transcript levels were depleted down to 9% of those of solvent-treated GI-ME-N controls and to 30% in CLB-GA cells (**Fig. 1a**). To establish whether *TERT* suppression is a general property of HDAC inhibition or restricted to specific isozyme classes or compounds, GI-ME-N and CLB-GA cells were exposed to a panel of pan-HDAC and isoform-selective inhibitors (**Fig. 1b**). Treatment with the alternative pan-HDAC inhibitors valproic acid and vorinostat reduced *TERT* transcript levels by up to 77% in GI-ME-N and up to 50% in CLB-GA cells. Similarly, treatment with entinostat (inhibiting HDACs 1–3 and 9) or mocetinostat (inhibiting HDACs 1–3 and 11) caused an up to 73% downregulation of *TERT* in GI-ME-N and 47% in CLB-GA cells, thus mirroring panobinostat treatment. Target discrimination using the HDAC1/2-selective inhibitor romidepsin suppressed *TERT* as effectively as broad-spectrum pan-HDAC inhibitors. Selective inhibition of HDAC2 with Santacruzamate A significantly reduced *TERT* levels to ∼60% in GI-ME-N and 76% in CLB-GA cells, relative to controls. In contrast, neither the HDAC8-selective inhibitor compound 2 nor bufexamac (inhibiting HDAC6 and HDAC10) altered *TERT* expression (**Fig. 1b**). To confirm the specific causative roles of individual class I HDAC isozymes, each of the eleven classical HDACs (HDACs 1–11) was systematically depleted using RNA interference in GI-ME-N cells. Specific knockdown of HDAC1 or HDAC2 individually suppressed *TERT* expression by approximately 30% in GI-ME-N cells (**Fig. 1c**), whereas siRNA-mediated depletion of HDACs 3-11 had no detectable effect (**Suppl. Fig. S2**). These results demonstrate that HDAC1 and HDAC2 are uniquely required to sustain *TERT* expression in *TERT*-rearranged neuroblastoma cells.

**Fig. 1.**
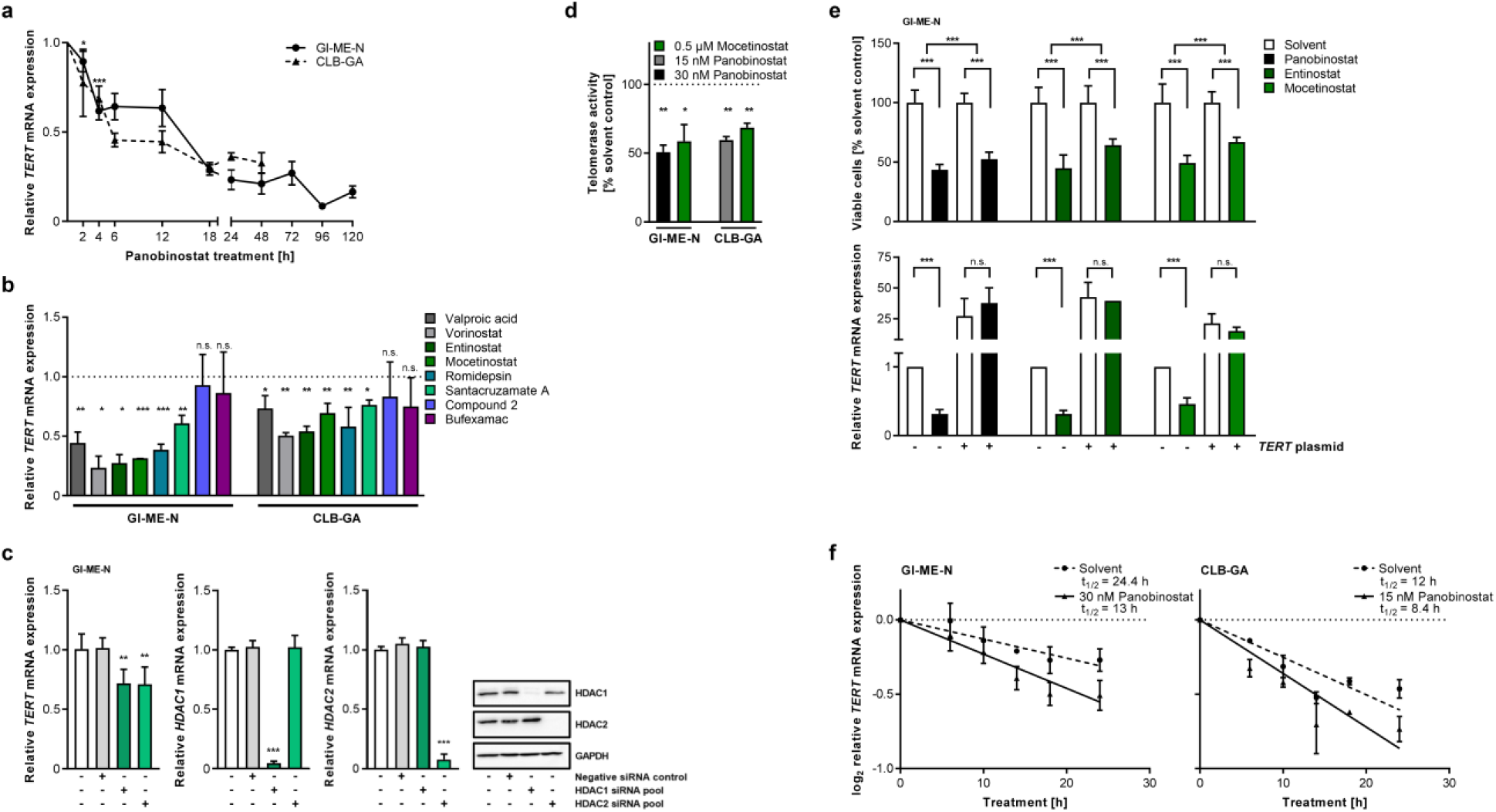
HDAC inhibition represses *TERT* expression, telomerase activity and *TERT* mRNA stability in *TERT*-rearranged neuroblastoma cells. **a**, Relative *TERT* mRNA expression measured by qRT-PCR in GI-ME-N and CLB-GA cells treated with panobinostat or solvent control (n≥3) over a 120 h time course. **b**, Relative *TERT* mRNA expression in GI-ME-N and CLB-GA cells treated with a panel of HDAC inhibitors or controls (n≥3). **c**, Influence of small interfering RNA (siRNA)-mediated knockdown (n≥3) of HDAC1 and HDAC2 on *TERT* mRNA expression in GI-ME-N cells. Knockdown efficacy was evaluated by qRT-PCR and Western blot analysis. **d**, Telomerase activity (% solvent control; n≥3) in GI-ME-N and CLB-GA cells treated with mocetinostat or panobinostat. **e**, Cell viability and relative *TERT* mRNA expression with (+) or without (-) TERT plasmid rescue at 72h of treatment with panobinostat, entinostat, mocetinostat or solvent control (n≥3). **f**, *TERT* mRNA decay kinetics and estimated transcript half-life (t_1/2_) in GI-ME-N and CLB-GA cells post-transcriptional block. Data (n≥3) represent mean ± SD and are shown on log_10_ scale. * *P* < 0.05; ** *P* < 0.01; *** *P* < 0.001.

To evaluate whether *TERT* transcript depletion functionally impairs the telomerase holoenzyme complex, enzymatic telomerase activity was quantified using a PCR-based ELISA assay^12^. Continuous treatment with panobinostat or mocetinostat over a 120 h time course reduced telomerase activity by approximately 2-fold in both GI-ME-N and CLB-GA cells (**Fig. 1d**). To demonstrate that tumor cell viability following HDAC inhibition is linked to *TERT* transcript level reduction, GI-ME-N cells were transiently transfected with a *TERT* cDNA expression plasmid in parallel to treatment with panobinostat, entinostat or mocetinostat. Ectopic re-expression of *TERT* cDNA significantly rescued cell viability, counteracting the loss of viability induced by panobinostat (by ∼9%), entinostat (by ∼19%), and mocetinostat (by ∼18%; **Fig. 1e**).

To define the molecular mechanism by which HDAC1/2 blockade reduces *TERT* transcript levels, we performed transcript decay kinetics following *de novo* transcriptional arrest with actinomycin D. Co-treatment of GI-ME-N and CLB-GA cells with actinomycin D and panobinostat markedly accelerated *TERT* mRNA decay compared to actinomycin D alone (**Fig. 1f**). Panobinostat reduced the estimated *TERT* transcript half-life (t_1/2_) from 24.4 h to 13.0 h in GI-ME-N cells (a 46.8% reduction in stability) and from 12.0 h to 8.4 h in CLB-GA cells (a 30.0% reduction; **Fig. 1f**). These kinetic data establish that HDAC1 and HDAC2 act post-transcriptionally to protect *TERT* mRNA from rapid degradation.

### HDAC inhibition does not alter local chromatin accessibility, active/repressive histone marks or DNA methylation at the *TERT*-rearranged breakpoint locus

To confirm that *TERT* downregulation is not caused by loss of local chromatin accessibility or transcriptional silencing at the translocation breakpoint, we first characterized the genomic architecture of the *TERT* rearrangement partner in GI-ME-N cells using long-read sequencing technologies (Oxford Nanopore and Pacific Biosciences). High-coverage long-read sequencing uniformly pinpointed a structural breakpoint within the 5′-untranslated region (5′-UTR) of *TERT* juxtaposed to a subtelomeric locus on chromosome 19q13.43 (SLC27A5/TAR1 region; **Suppl. Fig. S3a, b**), a rearrangement previously demonstrated^12^ on metaphase chromosomes by dual-color break-apart fluorescence in situ hybridization. We next performed chromatin immunoprecipitation coupled with high-throughput sequencing (ChIP-seq) in GI-ME-N cells treated with panobinostat (18 h) or solvent control to profile active (H3K4me3, H3K36me3, H3K27ac) and repressive (H3K9me3, H3K27me3) histone modifications across the breakpoint locus (**Suppl. Fig. S4a**). Comparative analysis of normalized ChIP-seq read counts revealed no significant treatment-induced changes in H3K4me3, H3K36me3, or H3K9me3 marks within the *TERT*-rearranged genomic region. Statistically significant increases in H3K27ac and H3K27me3 read counts were detected exclusively in genomic regions upstream of *TERT*, mapped specifically to the adjacent *ZBTB45* and *MZF1-AS1* gene loci rather than the *TERT* coding sequence or hijacked enhancer elements. Furthermore, array-based DNA methylation profiling using the Illumina Infinium Human MethylationEPIC BeadChip showed only a mild, non-specific global reduction in CpG methylation upon panobinostat exposure (**Suppl. Fig. S4b**) and no focal correlation with *TERT* transcript loss across the breakpoint locus (**Suppl. Fig. S4c**). Together, these epigenomic analyses demonstrate that the strong transcriptional drive provided by enhancer hijacking remains structurally intact during HDAC inhibition, confirming that HDAC1/2 regulate *TERT* through a post-transcriptional mechanism.

### Preventive panobinostat administration attenuates *TERT*-rearranged xenografted tumor growth

To determine whether the post-transcriptional dependency on HDAC1/2 translates into therapeutic efficacy *in vivo*, immunocompromised nude mice bearing subcutaneous *TERT*-rearranged xenografts were evaluated in preventive treatment settings. In GI-ME-N xenografts, daily intraperitoneal panobinostat administration attenuated tumor volume progression by 2.0-fold compared to solvent-treated controls over a 25-day monitoring period (**Fig. 2a**). In the CLB-GA xenograft model, preventive panobinostat treatment suppressed tumor growth by 3.7-fold over 29 days (**Fig. 2b**), resulting in a striking and statistically significant prolongation of overall survival (**Fig. 2b**, Log-rank *P* < 0.001). Histopathological assessment of harvested xenograft tissues confirmed >90% tumor cell content across GI-ME-N (**Suppl. Fig. S5a**) and CLB-GA preventive models (**Suppl. Fig. S5b**). Molecular analysis of harvested xenograft punch biopsies demonstrated robust intratumoral target repression (**Fig. 2c**). Quantitative RT-PCR confirmed that panobinostat treatment reduced intratumoral *TERT* mRNA transcript levels to 61% in GI-ME-N xenografts and to 56% in CLB-GA xenografts relative to vehicle controls (**Fig. 2c**). Concurrently, telomerase enzymatic activity in harvested tumor tissues was suppressed down to 10% in GI-ME-N xenografts and to 23% in CLB-GA xenografts (**Fig. 2c**), demonstrating effective *in vivo* modulation of the HDAC1/2-*TERT* axis.

**Fig. 2.**
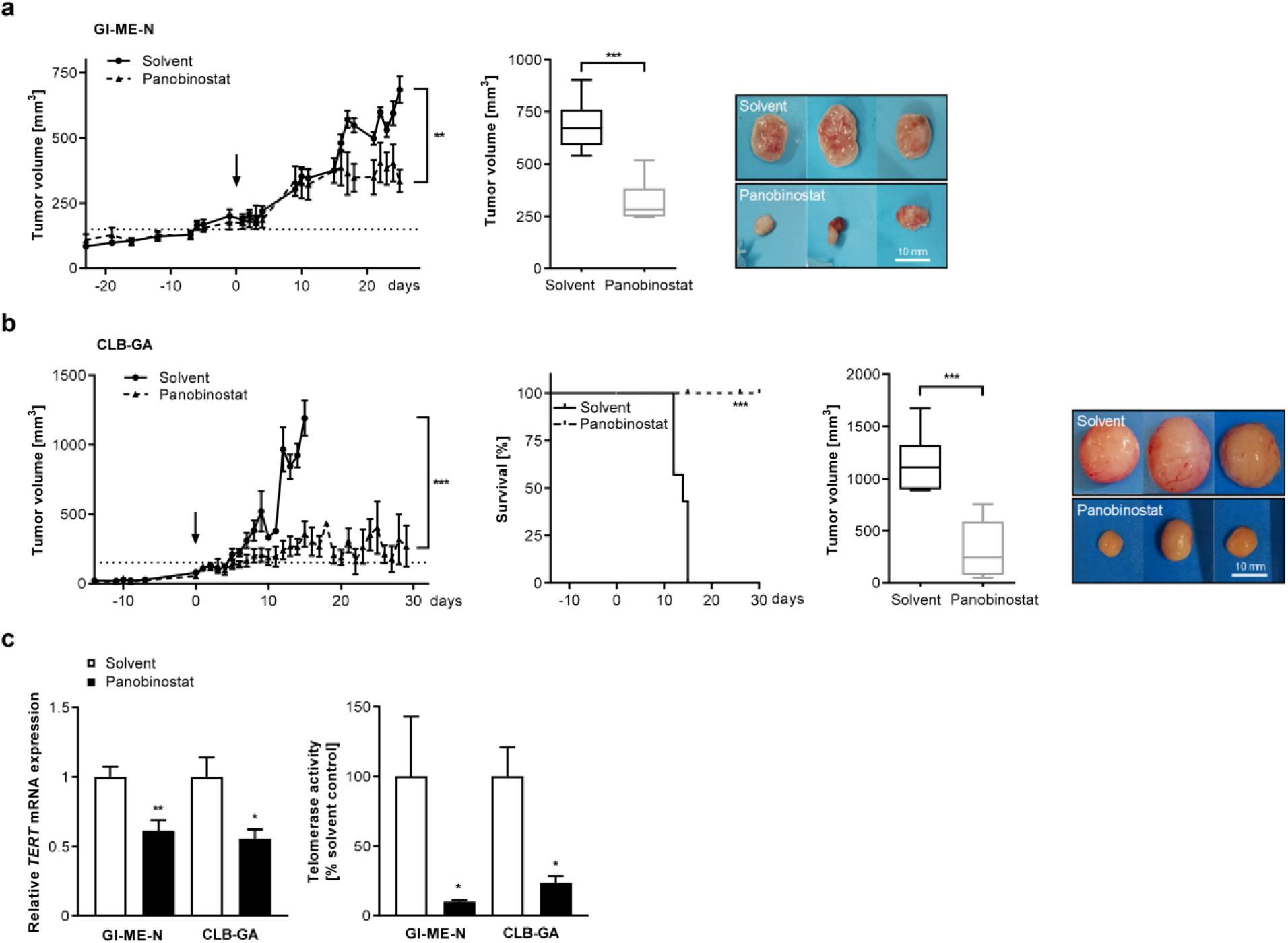
Panobinostat attenuates tumor growth, prolongs survival, and represses TERT in preventive xenograft models of *TERT*-rearranged neuroblastoma. **a**, Tumor volume progression, endpoint tumor volume box plot, and excised tumor photos from GI-ME-N preventive xenografts (n ≥ 6 per study group). **b**, Tumor volume progression, Kaplan-Meier survival curves, endpoint tumor volume, and excised tumor photos from CLB-GA preventive xenografts (n ≥ 6 per study group). **c**, Relative *TERT* mRNA expression and telomerase activity in harvested preventive xenograft tumor tissues (n ≥ 6 per study group). Data represent mean ± SD.* *P* < 0.05, ** *P* < 0.01, *** *P* < 0.001.

### Panobinostat therapy suppresses telomerase activity in established xenografts in mice

To mimic clinical presentation more closely, panobinostat efficacy was evaluated in an established therapeutic xenograft model using CLB-GA cells. Treatment was initiated only after tumors reached established palpable volumes (∼250-300 mm³). Therapeutic panobinostat administration arrested tumor progression over the 11-day treatment window, yielding a 6.7-fold reduction in final endpoint tumor volume compared to rapidly progressing vehicle-treated controls (**Fig. 3a**). Histological validation confirmed tumor cell-rich xenograft tissue (**Suppl. Fig. S5c**). Ex vivo molecular profiling of verified tumor biopsies revealed a 53% reduction in intratumoral *TERT* mRNA transcript levels (down to 47% of solvent controls; **Fig. 3b**) accompanied by a 78% inhibition of telomerase enzymatic activity (down to 22% of solvent controls; **Fig. 3b**). These results confirm that pharmacological HDAC inhibition effectively suppresses *TERT* and halts tumor growth in pre-established *TERT*-rearranged xenograft tumors.

**Fig. 3.**
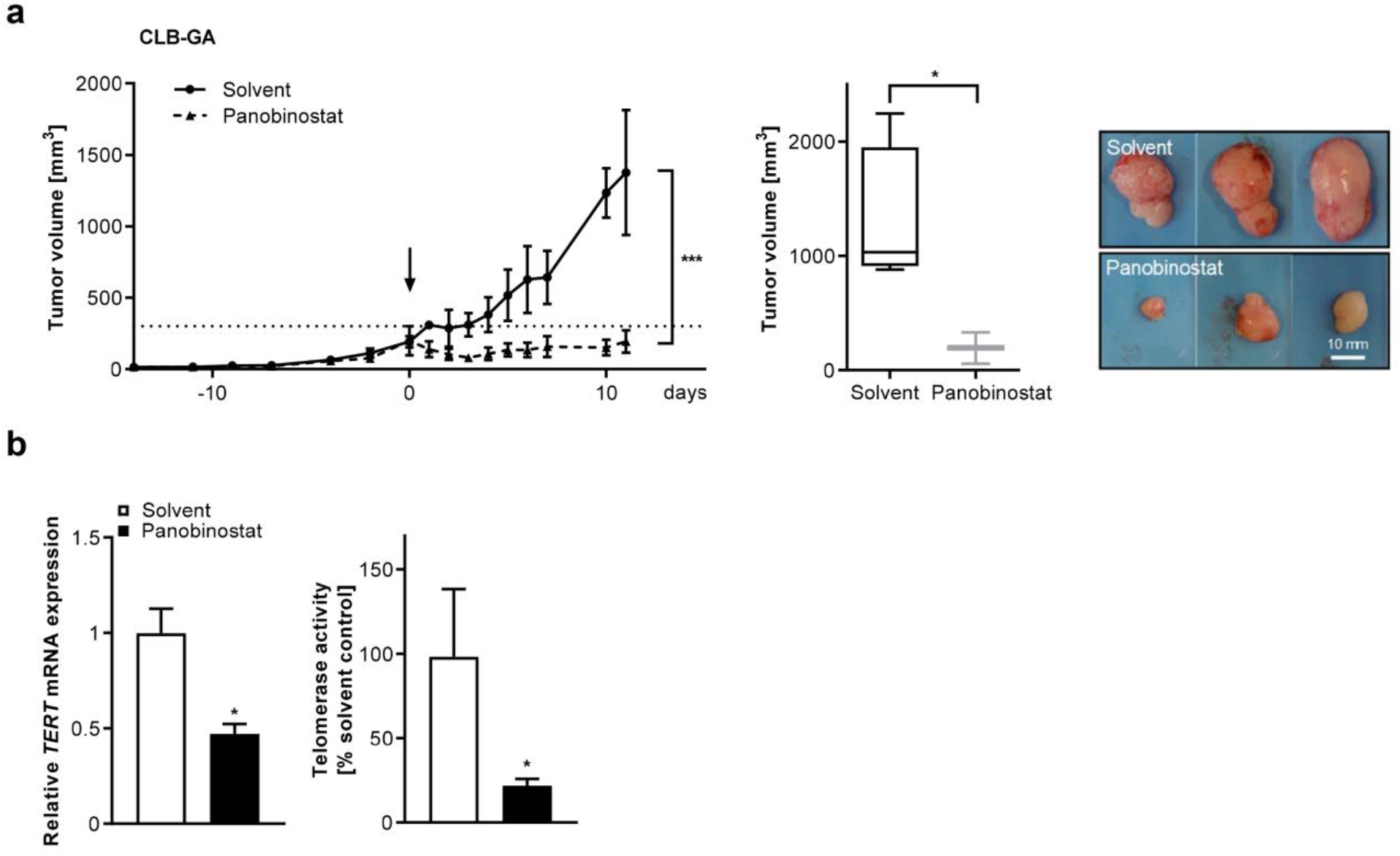
Therapeutic panobinostat administration suppresses established tumor growth and telomerase activity in *TERT*-rearranged neuroblastoma xenografts. **a**, Tumor volume progression, endpoint tumor volume box plot, and excised tumor photos from established CLB-GA therapeutic xenografts (n ≥ 3 per study group). **b**, Relative *TERT* mRNA expression and telomerase activity in harvested therapeutic xenograft tumor tissues (n ≥ 3 per study group). Data represent mean ± SD.* *P* < 0.05; *** *P* < 0.001.

### Panobinostat suppresses the *TERT*-associated 50-gene expression signature

High *TERT* expression in non-*MYCN*-amplified neuroblastoma is accompanied by the coordinated activation of a core 50-gene *TERT*-associated oncogenic expression signature^19^ encompassing essential cell cycle, mitotic, and DNA repair drivers. To assess whether post-transcriptional *TERT* transcript reduction dismantles this oncogenic network, we conducted gene set enrichment analysis (GSEA) across cell lines and xenograft models treated with panobinostat. GSEA dot plots and receiver operating characteristic (ROC) evidence plots demonstrated strong, highly statistically significant repression of the core 50-gene *TERT* signature across all contrast groups (**Fig. 4a, b**). Quantitative RT-PCR validation in CLB-GA cells exposed to panobinostat confirmed profound downregulation of representative signature hub genes including *EXO1, E2F8*, and *MIS18A* (**Fig. 4c, Suppl. Fig. S6a**). To prove that signature collapse is specifically mediated by *TERT* transcript depletion rather than non-specific off-target HDAC inhibitor toxicity, GI-ME-N cells transfected with a *TERT* cDNA expression plasmid were treated with panobinostat. Ectopic *TERT* re-expression significantly rescued and normalized mRNA levels of *EXO1, E2F8*, and *MIS18A* (**Fig. 4d**) as well as additional signature targets including *UHRF1*, *AURKA* and *BIRC5* (**Suppl. Fig. S6b**), showing a correlation between *TERT* transcript recovery and signature restoration. To extend beyond the targeted 50-gene TERT signature defined by Huang et al.^19^ and gain an unbiased, genome-wide view of the transcriptional networks regulated by panobinostat treatment, we performed genome-wide REACTOME pathway enrichment analysis based on the RNA-seq data obtained across all experimental models including CLB-GA and GI-ME-N cell lines, *in vivo* preventive xenograft tumors and a combined discovery cohort representing the merged dataset of cell lines and xenograft tumors. This unbiased analysis across all RNA-seq datasets revealed a widespread, highly concordant downregulation of core cellular programs (**Suppl. Fig. S7**), demonstrating that panobinostat treatment systematically suppresses broader biological processes intrinsically linked to TERT and telomere function. Beyond cell cycle progression, DNA replication and mitotic spindle assembly, REACTOME profiling specifically identified the significant enrichment and downregulation of pathways governing telomere maintenance, packaging of telomere ends, chromosome end preservation and DNA double-strand break repair across CLB-GA and GI-ME-N sample sets by panobinostat treatment (**Suppl. Fig. S7**). These findings suggest that post-transcriptional TERT mRNA destabilization by HDAC1/2 inhibition leads to the comprehensive shutdown of the broader genome-wide transcriptional program required to maintain telomere integrity and oncogenic proliferation in *TERT*-rearranged neuroblastoma.

**Fig. 4.**
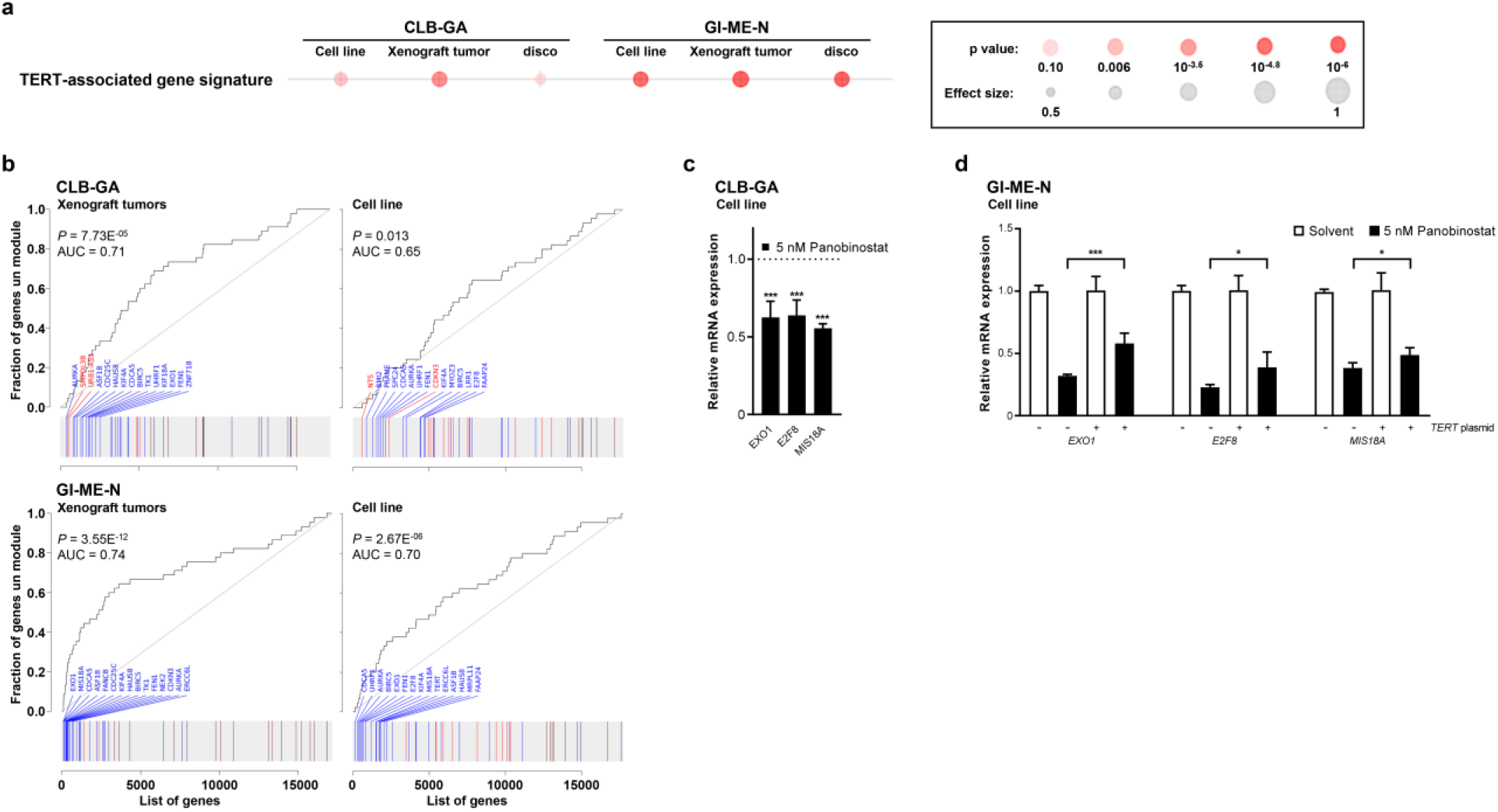
Panobinostat represses the *TERT*-associated 50-gene expression signature *in vitro* and *in vivo*, partially rescued by TERT re-expression. **a**, GSEA dot plot depicting statistical significance and effect size for the TERT-associated gene signature across contrasts. **b**, ROC curve plots for the TERT signature in CLB-GA and GI-ME-N xenografts and cell lines. **c**, qRT-PCR validation of representative TERT signature genes (EXO1, E2F8, MIS18A) in CLB-GA cells treated with panobinostat. **d**, qRT-PCR analysis of EXO1, E2F8, and MIS18A expression in GI-ME-N cells with (+) or without (-) TERT plasmid rescue. Data represent mean ± SD. * *P* < 0.05; *** *P* < 0.001.

In summary, our experimental findings establish a novel post-transcriptional mechanism in *TERT*-rearranged neuroblastoma (**Fig. 5**). While structural genomic rearrangements supply a strong transcriptional drive via enhancer hijacking, HDAC1 and HDAC2 serve as essential post-transcriptional guardians that actively protect *TERT* mRNA from rapid degradation. Pharmacological inhibition or depletion of HDAC1/2 accelerates *TERT* mRNA decay, reduces telomerase activity and disrupts the core 50-gene *TERT*-associated oncogenic network, driving antitumoral responses *in vitro* and *in vivo* (**Fig. 5**).

**Fig. 5.**
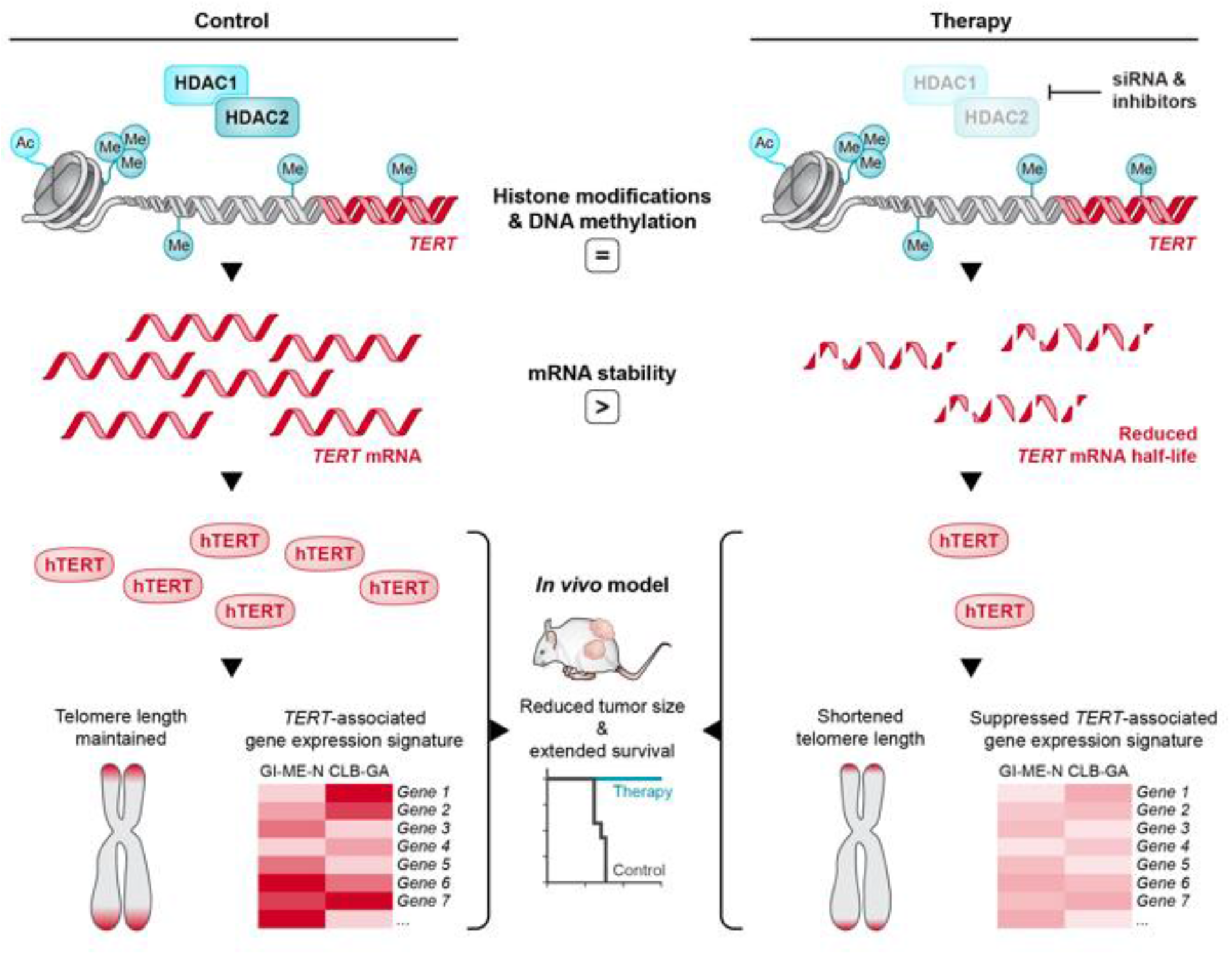
Graphical abstract summarizing the posttranscriptional regulation of *TERT* mRNA transcripts in *TERT*-rearranged neuroblastoma and indicating the experimental strategies applied.

## DISCUSSION

This study demonstrates that histone deacetylases 1 and 2 (HDAC1/2) act as essential post-transcriptional regulators that stabilize *TERT* mRNA transcripts in *TERT*-rearranged neuroblastoma cells. Pharmacologic inhibition of HDAC1/2 using the clinically approved pan-HDAC inhibitor panobinostat accelerates *TERT* mRNA transcript decay, reduces enzymatic telomerase activity, dismantles the core 50-gene TERT-associated oncogenic expression network^19^ and contributes to potent antitumoral responses in preclinical disease models. Genomic rearrangements at the *TERT* locus (5p15.33) represent a defining genetic driver in approximately 20–30% of high-risk neuroblastomas, demarcating a patient subgroup characterized by aggressive disease biology, high frequency of chemo-resistant relapse and dismal overall survival.^9,12,13^ These structural alterations are mutually exclusive with *MYCN* amplifications and *ATRX* loss-of-function mutations, establishing *TERT* activation as one of three primary, non-overlapping telomere maintenance mechanisms that confer cellular immortality in high-risk pediatric neuroblastoma.^9,12^ While direct pharmacological inhibitors of the catalytic telomerase reverse transcriptase subunit (TERT) have remained clinically unavailable so far, our study uncovers an actionable epigenetic vulnerability in this aggressive tumor subgroup.

The role of histone deacetylases in TERT regulation presents a well-documented epigenetic paradox across cell types and developmental states. In normal human somatic cells and telomerase-negative models, *TERT* is epigenetically silenced at the promoter level, where HDAC1 and HDAC2 serve as core catalytic subunits of repressive corepressor complexes, including Sin3B, NuRD, and CoREST.^25–27^ In these non-malignant contexts, pharmacological HDAC inhibition derepresses the *TERT* promoter, resulting in increased histone acetylation and transcriptional induction of *TERT* mRNA.^25,26^ Conversely, in *TERT*-rearranged neuroblastoma, structural genomic alterations supply a powerful, active transcriptional drive via enhancer hijacking that bypasses canonical promoter silencing mechanisms.^12,13^ Our ChIP-seq and DNA methylation profiling across the translocation breakpoint locus demonstrate that HDAC inhibition by panobinostat does not alter local chromatin accessibility, active/repressive histone marks (H3K27ac, H3K4me3, H3K9me3) or CpG methylation. Instead, our kinetic and functional data reveal a novel post-transcriptional dependency wherein HDAC1 and HDAC2 act as essential guardians that protect *TERT* mRNA transcripts from rapid degradation. This post-transcriptional mode of regulation distinctively differs from previously described post-translational mechanisms. For instance, in vascular smooth muscle cells, HDAC inhibition was reported to induce *TERT* transcription while simultaneously accelerating TERT protein degradation.^28^ While earlier reports in prostate cancer^29^ and hematological malignancies^30^ noted that HDAC inhibitors could reduce TERT transcript levels, the precise molecular mechanism remained unresolved. Our findings establish that in tumors fueled by enhancer hijacking, HDAC1 and HDAC2 are co-opted at the post-transcriptional level to preserve steady-state TERT mRNA levels, establishing an actionable RNA-level vulnerability in high-risk neuroblastoma.

Within the oncologic landscape, TERT activation is a ubiquitous hallmark across multiple human cancers but the dominant genetic and epigenetic mechanisms governing TERT deregulation vary markedly across tumor entities.^18,31^ In adult solid malignancies, TERT re-activation often occurs through highly recurrent, somatic single-nucleotide mutations in the *TERT* promoter region at positions -124 and -146 (designated C228T and C250T), which create *de novo* binding motifs for E26 transformation-specific family transcription factors.^32^ *TERT* promoter mutations are driving alterations in 40% to 70% in melanoma^33,34^, bladder^35^, liver^36^, and central nervous system cancers^37^, such as glioblastoma^38^. Pan-cancer genomic analyses encompassing over 1,200 whole-genome sequenced tumors across more than 20 cancer types demonstrated that structural variants located 0-100 kb upstream of *TERT* represent a second global, high-impact mechanism driving TERT overexpression across diverse cancer types including renal cell carcinoma, hepatocellular carcinoma, B-cell hematological malignancies and soft tissue sarcomas.^18,39^ Across these cancers, structural rearrangements reposition strong, active distal enhancers into proximity to the *TERT* coding sequence - a phenomenon termed enhancer hijacking - leading to epigenetic remodeling, high-level *TERT* transcription and elevated telomerase activity.^18^ In neuroblastoma, *TERT* promoter hotspot mutations are exceedingly rare. Instead, structural genomic rearrangements supply the primary transcriptional drive via enhancer hijacking.^12,13^ Our epigenomic profiling across the translocation breakpoint locus in *TERT*-rearranged GI-ME-N cells revealed that panobinostat treatment does not alter local chromatin accessibility, active/repressive histone modification marks (H3K27ac, H3K4me3, H3K9me3, H3K27me3, H3K36me3) or locus-wide DNA methylation beta-values. Instead, our functional genetic and pharmacological data establish a novel post-transcriptional mechanism. Class I HDACs – specifically HDAC1 and HDAC2 – are required post-transcriptionally to protect *TERT* mRNA from degradation. Specific siRNA-mediated knockdown of HDAC1 or HDAC2 reduced *TERT* mRNA transcript levels. Actinomycin D transcript decay kinetics confirmed that HDAC inhibition significantly impairs *TERT* mRNA stability, reducing the transcript half-life by 46.8% in GI-ME-N cells and by 30% in CLB-GA cells. Thus, while chromosomal rearrangements provide the initial enhancer-hijacking transcriptional boost, HDAC1 and HDAC2 serve as essential post-transcriptional guardians that prevent *TERT* mRNA decay, maintaining the elevated steady-state transcript levels necessary to sustain telomere lengthening and oncogenic signaling. Crucially, panobinostat was deployed in this study strictly as a chemical tool drug to unmask this epigenetic dependency, rather than as a proposed clinical monotherapy or combination therapy component. Uncovering that HDAC1/2 act as post-transcriptional guardians shifts the translational focus toward precision strategies that disrupt HDAC1/2-mediated RNA stabilization. Previous work by Chen and colleagues^40^ established that BET bromodomain inhibitors (e.g., OTX015) evict the epigenetic reader and co-activator BRD4 from hijacked enhancers to suppress *TERT* transcription and induce endoplasmic reticulum stress. Here, we expand upon this paradigm by uncovering a post-transcriptional layer of epigenetic regulation. This distinction between upstream transcriptional elongation and downstream post-transcriptional transcript preservation carries potential therapeutic implications. Because BET bromodomain inhibitors halt de novo transcript synthesis at hijacked super-enhancers, combining transcriptional blockade with post-transcriptional *TERT* transcript destabilization may provide a compelling dual-layer epigenetic strategy to completely inhibit TERT expression in *TERT* rearranged high-risk neuroblastoma.

A major functional consequence of high *TERT* expression in non-*MYCN*-amplified neuroblastoma is the coordinated activation of a core 50-gene TERT-associated expression signature.^19^ Originally identified by GSEA transcriptomic analysis of 498 patient tumors (GSE62564), this signature encompasses key oncogenic drivers and cell cycle regulators including BIRC5, AURKA, UHRF1, FEN1, E2F8, EXO1, MIS18A, CDCA5, ERCC6L, and ASF1B. Many genes overlap with the MYC and E2F transcriptional networks and are regulated by the super-enhancer co-activator Brd4 and cyclin-dependent kinases.^19^ In our study, panobinostat treatment strongly repressed this 50-gene signature across both cell lines and the xenograft models, causing marked downregulation of top signature components (*EXO1*, *E2F8*, *MIS18A*, *BIRC5*, *AURKA*, *UHRF1*). Ectopic plasmid mediated re-expression of *TERT* cDNA partially rescued cellular viability and normalized expression of key signature genes (*EXO1*, *E2F8*, *MIS18A*) despite continuous exposure to panobinostat. This genetic rescue confirms that panobinostat mediated suppression of this oncogenic network is in part driven by *TERT* mRNA transcript depletion. Signature components like BIRC5 and AURKA are therapeutic targets in high-risk neuroblastoma. Indirectly reducing their expression via HDAC1/2-mediated TERT mRNA destabilization offers a strategy to dismantle multiple oncogenic pathways simultaneously.

Translating these mechanistic insights *in vivo*, panobinostat demonstrated robust therapeutic efficacy in immunocompromised xenograft mouse models of *TERT*-rearranged neuroblastoma. In preventive treatment regimens, panobinostat significantly attenuated tumor growth, decreased final tumor volume and strongly prolonged overall survival of mice carrying CLB-GA xenografts. In established, therapeutic CLB-GA xenografts, panobinostat arrested tumor progression accompanied by a marked suppression of intratumoral *TERT* mRNA transcript levels and telomerase activity. These *in vivo* findings align with broader clinical frameworks established by Ackermann et al.^9^, which emphasize that aggressive high-risk neuroblastomas often combine telomere maintenance mechanisms (*TERT* rearrangements or *MYCN* amplifications) with co-occurring mutations in the RAS/MAPK/ALK pathway. In agreement with this multi-hit model, combining HDAC1/2 targeting drugs with further targeted agents such as the third-generation ALK inhibitor lorlatinib or the mTORC1 inhibitor rapamycin^41^ may be beneficial.

The clinical translation of anti-HDAC1/2 directed therapeutic regimens in *TERT*-driven neuroblastoma can be powerfully integrated with emerging liquid biopsy diagnostic platforms. Recent work by Hollander et al.^42^ demonstrated that droplet digital PCR quantification of patient-specific *TERT* rearrangement breakpoints in cell-free DNA isolated from peripheral blood and bone marrow plasma provides a sensitive, real-time biomarker for minimal residual disease monitoring. Cell-free tumor DNA *TERT* breakpoint dynamics captured therapy response, molecular remission and early relapse months before standard clinical imaging or conventional bone marrow cytology in individual patients.^42^ Integrating real-time ctDNA TERT breakpoint monitoring into clinical trial protocols may enable precise tracking of treatment response and early detection of resistance during HDAC inhibitor-containing therapeutic regimens.

While our study provides compelling preclinical evidence for targeting *TERT*-rearranged neuroblastoma via HDAC inhibition, several limitations warrant consideration. First, a major limitation in preclinical *TERT* rearrangement research is model availability. Only few cell line models harboring confirmed endogenous *TERT* rearrangements currently exist^12^, inherently restricting the breadth of *in vitro* cell models available for functional validation. Second, our *in vivo* xenograft studies were conducted using subcutaneous cell line-derived xenografts in immunocompromised mice. While these models demonstrated clear antitumoral efficacy, they lack an intact adaptive immune system and do not fully recapitulate the complex human neural crest tumor microenvironment or orthotopic stromal interactions. Third, although we established that HDAC1 and HDAC2 depletion accelerates *TERT* mRNA transcript decay, the exact post-transcriptional machinery such as specific RNA-binding proteins, m6A epitranscriptomic modifications or deadenylase complexes directly deacetylated by HDAC1/2 to stabilize TERT mRNA remains to be fully dissected in future biochemical studies.

Although small-molecule HDAC inhibitors have demonstrated preclinical efficacy, their clinical utility in solid tumors remains severely constrained by occupancy-driven limitations, narrow therapeutic windows and broad dose-limiting toxicities such as severe thrombocytopenia, fatigue, and gastrointestinal distress.^43^ Targeted protein degradation using proteolysis targeting chimeras (PROTACs) circumvents these barriers through sub-stoichiometric, event-driven catalysis that selectively destroys target proteins rather than relying on active-site occupancy.^44,45^ By inducing the proteasomal degradation of HDAC1 and HDAC2, PROTACs eliminate both catalytic deacetylase activity and non-enzymatic scaffolding within nuclear co-repressor complexes.^46^ In *TERT*-rearranged neuroblastoma, where HDAC1 and HDAC2 act as essential post-transcriptional guardians of *TERT* transcript stability, selective HDAC1/2 degraders may offer a translationally superior avenue to accelerate *TERT* mRNA decay, abolish telomerase activity and collapse the core 50-gene TERT-associated oncogenic network^19^ at lower systemic drug exposures. Sparing non-target Class II/IV HDACs is likely to minimize off-target toxicities, providing a precision-medicine framework to target epigenetic dependencies in high-risk neuroblastoma driven by TERT rearrangements.

In conclusion, our findings establish that the class I HDAC family members HDAC1 and HDAC2 act as essential post-transcriptional guardians of *TERT* mRNA transcript stability in *TERT*-rearranged high-risk neuroblastoma. Pharmacological inhibition of HDAC1/2 using clinically approved agents such as panobinostat destabilizes *TERT* mRNA, suppresses telomerase enzymatic activity and inhibits the downstream *TERT*-associated oncogenic network. This post-transcriptional targetability deserves further evaluation for precision medicine approaches tailored to suppress the malignant machinery fueled by genomic *TERT* rearrangements in high-risk neuroblastoma.

## MATERIALS AND METHODS

### Drug testing in subcutaneous neuroblastoma xenografts in mice

Viable GI-ME-N (5 x 10^6^) or CLB-GA (2 x 10^7^) cells were suspended in 100 µl Matrigel™ (Corning Life Sciences, Tewksbury, MA, USA) and subcutaneously implanted in the flanks of 6-week-old female NMRI-Foxn1^nu/nu^ mice. Mice were randomly assigned to either panobinostat (Selleck Chemicals, Houston, U.S.) or control groups after tumors reached 100-150 mm^3^ (for preventive testing settings, n ≥ 6 mice per study group) or 250-300 mm^3^ (for therapeutic testing settings, n ≥ 3 mice per study group). Mice were injected intraperitoneally with 15 mg/kg/d panobinostat (in 5% glucose) or vehicle alone (control) once daily for 5 consecutive days followed by 2 days off for up to 30 d or until tumor size exceeded 1500 mm³, when mice were euthanized by cervical dislocation. Tumor size was measured daily by caliper, and volume was calculated using V=π/6 (d1 x d2 x d3), where d1 was the largest and d3 the smallest tumor diameter. Xenograft tumor histology was examined in formalin-fixed, paraffin-embedded sections stained with hematoxylin and eosin. Fresh tumor tissue was snap frozen in liquid nitrogen for molecular analyses. All experiments were performed in accordance with the “Guidelines for accommodation and care of laboratory animals” by the Council of Europe, and the protocol approved by the local ethics commission for animal experimentation (Charité # G 0173/17).

### Cell culture and *in vitro* treatment

The neuroblastoma cell lines, GI-ME-N (RRID: CVCL_1232, obtained from the DSMZ, Braunschweig, Germany) and CLB-GA (RRID: CVCL_9529) were maintained at 37°C and 5% CO_2_ without antibiotics and supplemented with 10% fetal calf serum (Merck, Darmstadt, Germany) and 1% non-essential amino acids (Biochrom, Berlin, Germany). Medium base (Life Technologies, Karlsruhe, Germany) was DMEM for the CLB-GA line and RPMI for GI-ME-N. Cell lines were authenticated using high-throughput SNP-based assays and regularly monitored for mycoplasma using PlasmoTest™ (InvivoGen) according to the manufacturer’s instructions. Viable cells were semi-automatically counted on a VI-CELL-XR Cell Viability Analyzer (Beckman Coulter, Brea, CA, U.S.) using the Trypan blue exclusion method. HDAC inhibitors were prepared as stock solutions (**Suppl. Table S1**) in dimethyl sulfoxide (DMSO) or phosphate-buffered saline (valproic acid). Final DMSO concentration in medium was <0.1%. RNA synthesis was inhibited by a final concentration of 4 µM actinomycin D (Sigma-Aldrich, Steinheim, Germany) in medium. For gene expression knockdown, single short interfering RNAs (siRNAs) or pools (20 nM total concentration) consisting of 30 different siRNAs (**Suppl. Table S2**) were transiently transfected into cells using Lipofectamine™ RNAiMAX (Thermo Fisher Scientific, Waltham, MA, USA) according to the manufacturer’s directions. For plasmid transfection, Effectene^TM^ (Qiagen) was used as the transfection reagent according to the manufacturer’s manual. The pBABE-neo-hTERT plasmid was a gift from Bob Weinberg^47^ (Addgene plasmid 1774), as well as the empty vector pBABE-neo (Addgene plasmid 1767), serving as control.

### ChIP-seq analysis of histone modifications

Cells were lysed for chromatin immunoprecipitation (ChIP) in buffer containing 50 mM Tris-HCl (pH 8.1), 1 % SDS, 10 mM EDTA and cOmplete^®^ protease inhibitor cocktail (Roche), and sonicated (Covaris Inc, Woburn, MA, USA) to produce chromatin fragments 200-800 bp in length before use in the ChIP Assay Kit (Merck Millipore, Darmstadt, Germany) according to the manufacturer’s instructions with antibodies listed in **Suppl. Table S3**. Libraries were prepared using the NEBNext^®^ ChIP-Seq Library Prep Reagent Set (Illumina, San Diego, CA, U.S.) according to the manufacturer’s protocol with a step selecting for fragments of approximately 300 bp length. Pooled libraries were sequenced on the Illumina HiSeq 4000 sequencing platform in 50 bp single-read mode. Sequencing reads were aligned to the GRCh38 reference human genome for peak discovery and annotation to functional gene regions before assessing probe occupancy (details in the **Supplementary Information**).

### DNA methylation profiling

Samples (350-500 ng DNA) were bisulfite converted with the EZ-96 DNA Methylation Kit (Zymo Research, Freiburg, Germany) according to the manufacturer’s instructions before amplification, fragmentation (300-600 bp) and microarray hybridization (Illumina Infinium Human MethylationEPIC Beadchip) using a standard Illumina protocol. Microarrays were scanned in the Illumina iScan-Scanner using their standard scanning protocol. Data files were processed (details in **Supplementary Information**) using the minfi R package^48^ before probe-wise differential methylation analysis and prediction of differentially methylated regions (details in **Supplementary Information**).

### Expression analysis

To compare treatment effects in the GI-ME-N and CLB-GA cell lines, libraries were prepared from samples (500 ng total RNA input) using the Illumina TruSeq (Illumina, San Diego, CA, U.S.) mRNA stranded protocol for multiplexed RNA sequencing (6 libraries/lane) on a Hiseq 4000 in 100 paired-end mode. A minimum of 48 million read pairs were obtained per library that were mapped to the GRCh38 reference human genome using STAR software^49^ (v 2.7.5a), and gene-level quantification was performed employing GENCODE (version 25) gene annotations for use in differential expression analysis and gene set enrichment for signatures in the Molecular Signatures Database^50^, KEGG pathways and the custom 50-gene, “*TERT-*associated gene expression signature”^19^ (details for data analysis in the **Supplementary Information**).

Total RNA isolated from cell lines or tissues using the RNeasy Mini Kit (Qiagen, Hilden, Germany) was converted to cDNA for quantitative reverse-transcriptase PCR (qRT-PCR) using the First-Strand cDNA Synthesis Kit (Thermo Fisher Scientific). Relative *TERT* expression was measured using SYBR Green (Roche, Basel, Switzerland) and specific primer pairs (**Suppl. Table S4**) in the StepOnePlus Real-time PCR system (Applied Biosystems, Foster City, CA, USA), and normalized to the geometric mean of *RPS18*, *HPRT* and *SDHA* expression.^51^ *HPRT* and *SDHA* are consistently expressed in INRG stage M and MS neuroblastomas^52^ and can be used as housekeeping genes. Data were analyzed using version 2.3 StepOne™ software. Expression changes in cell lines were calculated using the ΔΔC_t_ method, and a standard curve was used for xenograft tumor analyses.

Cells were lysed for western blotting using antibodies (**Suppl. Table S5**) to detect HDAC1 and 2 in buffer containing 20 mM Tris-HCl, 7 M urea, 0.01 % Triton X-100, 100 mM DTT, 40 mM MgCl_2_ and cOmplete^®^ protease inhibitor cocktail. Proteins were visualized using the Western Lightning Plus-ECL detection solution (Perkin Elmer, USA) and the Quantum ST5 imaging system (Vilber Lourmat, France). Ratios were calculated with the Bio-1D Version 12.10a software (Peqlab, Erlangen, Germany).

### Telomeric repeat amplification assay

Telomerase activity was assessed in extracts from cell cultures or xenograft tumor tissues using the TeloTAGGG™ Telomerase PCR ELISA PLUS assay (Roche, Rotkreuz, Switzerland) according to the manufacturer’s instructions.

### Statistical analyses

Differences between treatment cohorts in mouse xenograft experiments were determined using area under the curve and Kaplan-Meier survival analyses with log-rank statistics. Comparisons between transcript and telomerase activity levels were conducted using Student’s unpaired two-sided and one-sample t-tests. Analyses were conducted using the GraphPad Prism version 7.0 (GraphPad Software, La Jolla, CA, U.S.).

## ACKNOWLEDGEMENTS

We thank Jasmin Wünschel, Daniela Tiburtius and Jeannette Mothes for excellent technical assistance, Nicole Hübener for project management, Kathy Astrahantseff for critical discussions and Sandy Westermann for editing of the graphical abstract. We thank Förderverein KINDerLEBEN e.V. for supporting A. Szymansky and S. Fuchs through seed project funding and the DKFZ High Throughput Sequencing Unit for providing services.

## Data availability

ChIP-seq data, RNA-seq data and DNA methylation profiling data generated in this study have been deposited in the NCBI Gene Expression Omnibus (GEO) under SuperSeries accession numbers GSE155705 (ChIP-seq), GSE155707 (RNA-seq) and GSE155703 (methylation). Pacific Biosciences (PacBio) and Oxford Nanopore long-read sequencing data are available through the NCBI Sequence Read Archive under accession number SRR12476600. Publicly available microarray and RNA-seq datasets analyzed in this study were retrieved from GEO (GSE49711) and the R2 platform (https://r2platform.com). Unprocessed source data and supplementary materials are provided with the paper or are available from the corresponding author upon request.

