## Supplementary material for "HDAC1/2 maintain *TERT* mRNA stability in *TERT*-rearranged neuroblastoma": Suppl Data

**Table of Contents**

**Supplementary Materials and Methods** starting p. 2

**Supplementary Tables** starting p. 6

**Supplementary Figures** starting p. 11

### SUPPLEMENTARY MATERIALS AND METHODS

#### Long-read sequencing

High molecular weight DNA was extracted from GI-ME-N cells using the MagAttract HMW DNA Kit (Qiagen) or the smart DNA prep kit (Jena Analytik, Jena, Germany). DNA content was measured with a Qubit 3.0 Fluorometer (Thermo Fisher), and sample quality was confirmed using an Agilent 4200 TapeStation (Santa Clara, CA, USA). Libraries were prepared for Nanopore long-read DNA sequencing using the Ligation Sequencing Kit (SQK-LSK109, Oxford Nanopore Technologies Ltd., Oxford, UK), and sequenced on a R9.4.1 MinION flowcell (FLO-MIN106, Oxford Nanopore Technologies Ltd.). Raw Nanopore sequencing signals (FAST5 format) were base-called using Guppy (v2.3.7; Oxford Nanopore Technologies Ltd.) under default parameters. Demultiplexing and adapter trimming were conducted using qcat (v1.0.1; Oxford Nanopore Technologies Ltd.). Read length distributions and quality metrics were evaluated using NanoPlot (v1.0.0) as previously described.[1] High-quality filtered long reads were mapped to the GRCh38 human reference genome using minimap2 (v2.16-r922) with default parameters for Nanopore genomic reads.

Libraries were prepared for PacBio long-read DNA sequencing (Pacific Biosciences PacBio Sequel II system) according to the manufacturer's protocol. Circular consensus sequencing (CCS/HiFi) reads were aligned to the GRCh38 human reference genome using pbmm2 (PacBio SMRT Link v9.0.0) under default parameter settings. Structural genomic variants were detected using pbsv (PacBio SMRT Link v9.0.0) following a two-step workflow. Structural variant signatures were first identified in discovery mode (pbsv discover), utilizing tandem repeat annotations (PacBio_tandem_repeats.bed) with thresholds set for minimum signature length (--min-svsig-length 1000) and mapping quality (--min-mapq 20). Structural variant genotypes across candidate loci were subsequently called using calling mode with parameter settings: call-min-reads-all-samples, 2; call-min-reads-one-sample, 2; call-min-reads-per-strand-all-samples, 0; call-min-read-perc-one-sample, 10; call-min-bnd-reads-all-samples, 2; min-sv-length, 1000). Variant calls overlapping blacklisted ENCODE regions were filtered out.

#### Translocation breakpoint mapping and de novo assembly

Structural rearrangement breakpoints at the *TERT* 5′ region (within 100 bp of the transcription start site) were identified by inspecting split reads and supplementary alignments in the Integrative Genomics Viewer (IGV, v2.3.93). To resolve the exact junction sequence at single-nucleotide resolution, candidate long reads spanning the 5′ TERT locus were assembled de novo. All-against-all read overlap mapping was performed using minimap2 (v2.17-r941; preset -x ava-ont), followed by string-graph assembly with miniasm (v0.3-r179; flags -1 -2 -c 2 -e 2 -f) and consensus polishing using minipolish (v0.1.3). The assembled contigs were evaluated by re-mapping raw sequencing reads alongside reference genomic regions flanking the translocation breakpoint (GRCh38: chr5:1,250,000–1,300,000 and chr19:58,570,000–58,610,000) using minimap2 (preset -ax map-ont). Nucleotide-level alignments across the breakpoint junction were visually validated using samtools tview.

**RNA-sequencing and differential expression analysis**

Differential gene expression analysis was conducted using the DESeq2 Bioconductor package (v1.26).[2] Low-abundance transcripts were excluded prior to statistical evaluation by retaining only genes with at least 10 total counts across all libraries and a minimum of 5 counts in at least 3 samples. Built-in independent filtering was utilized within DESeq2 to maximize testing power, and raw p-values were adjusted for multiple testing using the Benjamini–Hochberg false discovery rate (FDR) procedure.[3]

**Gene set enrichment analysis**

Gene set enrichment analysis was performed using the CERNO algorithm in the tmod R package (v0.46.1).[4] Pathway gene sets were retrieved from the Molecular Signatures Database (MSigDB)[5] using msigdbr (v7.1.1), focusing on REACTOME pathways and the custom 50-gene TERT-associated expression signature[6]. Overrepresentation and pathway enrichment of MSigDB gene sets and KEGG pathways in paired panobinostat- versus solvent-treated neuroblastoma cell datasets were evaluated using clusterProfiler (v3.14.3).[7] To identify concordantly enriched pathways across both cell lines and xenograft mouse models, the disco scoring framework was applied to differential expression profiles using tmod as previously described.[8] Joint enrichment testing encompassed four pairwise contrasts (panobinostat vs. solvent) and two disco concordance comparisons. Gene sets were filtered to exclude those with extreme sizes (>80 or <10 genes) or those failing to meet significance thresholds (AUC > 0.7 and false discovery rate q < 0.001 in at least one comparison per database). To remove redundant pathways, candidate gene sets were ordered by their combined p-values across comparisons, and pairwise Jaccard similarity indices were calculated sequentially; gene sets with a Jaccard index >0.75 were hierarchically pruned. GSEA profiles were visualized using the evidencePlot and tmodPanelPlot utilities in tmod.

**ChIP-seq data analysis**

Single-read sequencing reads were aligned to the GRCh38 human reference genome using Rsubread.[9] High-quality, properly paired reads were retained, and genome-wide coverage tracks were generated using bamCoverage from deepTools.[10] Peak calling was performed using MACS2[11], with pipeline quality control evaluated using ChIPQC[12] Differential binding analysis was conducted separately for each histone modification using DiffBind with minOverlap=2, incorporating DESeq2 for differential occupancy testing. Peaks were assigned to genomic features - including promoters, transcriptional start site (TSS) regions (defined as

±3 kb from the TSS), and intergenic regions - using the annotatePeak function in ChIPseeker[13]. Track alignments and differential peaks were visually inspected using the Integrative Genomics Viewer (IGV).[14]

**Processing of bisulfite DNA sequencing files for DNA methylation data analysis**

DNA methylation array signal intensities were normalized across samples using quantile normalization.[15,16] Probes with detection *P*-values above 0.01 and probes recognized as unreliable due to cross-reactivity or underlying single-nucleotide variants were filtered out prior to downstream analysis.[17] Probes were annotated using the IlluminaHumanMethylationEPICanno.ilm10b4.hg19 Bioconductor package.[18] Probe-wise differential methylation analysis was performed using the limma R package[19] on M-values, whereas beta-values were utilized for data visualization and interpretation.[20] Differentially methylated regions (DMRs) were identified using the DMRcate package.[21]


1. De Coster, W., D'Hert, S., Schultz, D.T., Cruts, M. & Van Broeckhoven, C. NanoPack: visualizing and processing long-read sequencing data. *Bioinformatics* **34**, 2666–2669 (2018).
2. Love, M.I., Huber, W. & Anders, S. Moderated estimation of fold change and dispersion for RNA-seq data with DESeq2. *Genome Biol* **15**, 550 (2014).
3. Benjamini, Y. & Hochberg, Y. Controlling The False Discovery Rate - A Practical And Powerful Approach To Multiple Testing. *J. Royal Statist. Soc., Series B* **57**, 289–300 (1995).
4. Zyla, J. *et al.* Gene set enrichment for reproducible science: comparison of CERNO and eight other algorithms. *Bioinformatics* **35**, 5146–5154 (2019).
5. Liberzon, A. *et al.* The Molecular Signatures Database (MSigDB) hallmark gene set collection. *Cell Syst* **1**, 417–425 (2015).
6. Huang, M. *et al.* Epigenetic Targeting of TERT-Associated Gene Expression Signature in Human Neuroblastoma with TERT Overexpression. *Cancer Res* **80**, 1024–1035 (2020).
7. Yu, G., Wang, L.G., Han, Y. & He, Q.Y. clusterProfiler: an R package for comparing biological themes among gene clusters. *OMICS* **16**, 284–7 (2012).
8. Domaszewska, T. *et al.* Concordant and discordant gene expression patterns in mouse strains identify best-fit animal model for human tuberculosis. *Sci Rep* **7**, 12094 (2017).
9. Liao, Y., Smyth, G.K. & Shi, W. The R package Rsubread is easier, faster, cheaper and better for alignment and quantification of RNA sequencing reads. *Nucleic Acids Res* **47**, e47 (2019).
10. Ramirez, F., Dundar, F., Diehl, S., Gruning, B.A. & Manke, T. deepTools: a flexible platform for exploring deep-sequencing data. *Nucleic Acids Res* **42**, W187–91 (2014).
11. Feng, J., Liu, T., Qin, B., Zhang, Y. & Liu, X.S. Identifying ChIP-seq enrichment using MACS. *Nat Protoc* **7**, 1728–40 (2012).
12. Carroll, T.S., Liang, Z., Salama, R., Stark, R. & de Santiago, I. Impact of artifact removal on ChIP quality metrics in ChIP-seq and ChIP-exo data. *Front Genet* **5**, 75 (2014).
13. Yu, G., Wang, L.G. & He, Q.Y. ChIPseeker: an R/Bioconductor package for ChIP peak annotation, comparison and visualization. *Bioinformatics* **31**, 2382–3 (2015).
14. Thorvaldsdottir, H., Robinson, J.T. & Mesirov, J.P. Integrative Genomics Viewer (IGV): high-performance genomics data visualization and exploration. *Brief Bioinform* **14**, 178–92 (2013).
15. Aryee, M.J. *et al.* Minfi: a flexible and comprehensive Bioconductor package for the analysis of Infinium DNA methylation microarrays. *Bioinformatics* **30**, 1363–9 (2014).
16. Touleimat, N. & Tost, J. Complete pipeline for Infinium((R)) Human Methylation 450K BeadChip data processing using subset quantile normalization for accurate DNA methylation estimation. *Epigenomics* **4**, 325–41 (2012).
17. Pidsley, R. *et al.* Critical evaluation of the Illumina MethylationEPIC BeadChip microarray for whole-genome DNA methylation profiling. *Genome Biol* **17**, 208 (2016).
18. KD, H. IlluminaHumanMethylationEPICanno.ilm10b4.hg19: Annotation for Illumina's EPIC methylation arrays. doi:10.18129/B9.bioc.IlluminaHumanMethylationEPICanno.ilm10b4.hg19,

R package version 0.6.0, *https://bioconductor.org/packages/IlluminaHumanMethylation EPICanno.ilm10b4.hg19.*

19. Smyth, G.K. limma: Linear Models for Microarray Data. In: Gentleman R, Carey VJ, Huber W, Irizarry RA, Dudoit S (eds) Bioinformatics and Computational Biology Solutions Using R and Bioconductor Statistics for Biology and Health Springer, New York, NY 2005.
20. Du, P. *et al.* Comparison of Beta-value and M-value methods for quantifying methylation levels by microarray analysis. *BMC Bioinformatics* **11**, 587 (2010).
21. Peters, T.J. *et al.* De novo identification of differentially methylated regions in the human genome. *Epigenetics Chromatin* **8**, 6 (2015).
22. Szymansky, A. *et al.* Neuroblastoma Risk Assessment and Treatment Stratification with Hybrid Capture-Based Panel Sequencing. *J Pers Med* **11**(2021).

### SUPPLEMENTARY TABLES

#### Supplementary Table S1. HDAC inhibitors.

| Product | Supplier | Stock concentration | Inhibitory profile |
|---|---|---|---|
| Bufexamac | Selleck Chemicals[1] | 100 nM | HDAC6, HDAC10 |
| Compound 2 | Selleck Chemicals | 100 mM | HDAC8 |
| Entinostat | Selleck Chemicals | 1 mM | HDAC1-3, HDAC9 |
| Mocetinostat | Selleck Chemicals | 10 mM | HDAC1-3, HDAC11 |
| Panobinostat | Selleck Chemicals | 1 mM | HDAC classes I, II, IV |
| Romidepsin | Selleck Chemicals | 10 µM | HDAC1, HDAC2 |
| Santacruzamate A | Biozol[2] | 1 µM | HDAC2 |
| Vorinostat | Selleck Chemicals | 1 mM | HDAC classes I, II, IV |

[1] Houston, TX, U.S.A.
[2] Eching, Germany

### Supplementary Table S2. Small interfering RNAs.

| Product | Supplier |
|---|---|
| siPool directed against **HDAC1** (human)<br>Gene ID: 3065<br>Transcripts targeted: NM_004964, XM_011541309 | siTOOLs Biotech GmbH[1] |
| siPool directed against **HDAC2** (human)<br>Gene ID: 3066<br>Transcripts targeted: NM_001527, NR_033441, NR_073443, XM_011535788, XM_017010799 | siTOOLs Biotech GmbH |
| Standard negative control siPOOL | siTOOLs Biotech GmbH |
| Hs_**HDAC3**_4 FlexiTube siRNA<br>Hs_**HDAC3**_6 FlexiTube siRNA | Qiagen[2]<br>Qiagen |
| Hs_**HDAC4**_7 FlexiTube siRNA<br>**HDAC4** Silencer Pre-designed siRNA, #107926 | Qiagen<br>Thermo Fisher Scientific[3] |
| Hs_**HDAC5**_1 FlexiTube siRNA<br>Hs_**HDAC5**_4 FlexiTube siRNA | Qiagen<br>Qiagen |
| Hs_**HDAC6**_2 FlexiTube siRNA<br>Hs_**HDAC6**_10 FlexiTube siRNA | Qiagen<br>Qiagen |
| Hs_**HDAC7**_6 FlexiTube siRNA<br>Hs_**HDAC7**_10 FlexiTube siRNA | Qiagen<br>Qiagen |
| Hs_**HDAC8**_5 FlexiTube siRNA<br>Hs_**HDAC8**_6 FlexiTube siRNA | Qiagen<br>Qiagen |
| Hs_**HDAC9**_1 FlexiTube siRNA<br>Hs_**HDAC9**_4 FlexiTube siRNA | Qiagen<br>Qiagen |
| Hs_**HDAC10**_1 FlexiTube siRNA<br>Hs_**HDAC10**_4 FlexiTube siRNA | Qiagen<br>Qiagen |
| **HDAC11** Silencer Pre-designed siRNA, #130749<br>Hs_**HDAC11**_7 FlexiTube siRNA | Thermo Fisher Scientific<br>Qiagen |
| Allstars Negative Control siRNA<br>siGENOME RISC-Free Control siRNA | Qiagen<br>Thermo Fisher Scientific |

1 Planegg, Germany
2 Hilden, Germany
3 Waltham, MA, U.S.A.

### Supplementary Table S3. ChIP-grade antibodies and isotype control antibody.

| Target protein | Antibody source / Type | Supplier ID & Clone |
|---|---|---|
| Human H3K9Me3 | Rabbit polyclonal | Abcam[1], ab8898 |
| Human H3K4Me3 | Rabbit polyclonal | Abcam, ab8580 |
| Human H3K36Me3 | Rabbit polyclonal | Abcam, ab9050 |
| Human H3K27Me3 | Rabbit polyclonal | Active Motif[2], 39155 |
| Human H3K27Ac | Rabbit polyclonal | Abcam, ab4729 |
| | Rabbit IgG | Santa Cruz Biotechnology[3], sc-2027 |

1 Cambridge, UK
2 Carlsbad, CA, U.S.A.
3 Dallas, TX, U.S.A.

### Supplementary Table S4. Primers for quantitative RT-PCR.

| Gene | Supplier ID[1] | Forward primer | Reverse primer |
|---|---|---|---|
| *ASF1B* | QT00007224 | | |
| *AURKA* | QT00038213 | | |
| *BIRC5* | QT00081186 | | |
| *CDCA5* | QT00067179 | | |
| *E2F8* | QT00098294 | | |
| *ERCC6L* | QT01021601 | | |
| *EXO1* | QT00080717 | | |
| *FEN1* | QT00064722 | | |
| *HPRT* | | 5'- TGA CAC TGG CAA AAC AAT GCA -3' | 5'- GGT CCT TTT CAC CAG CAA GCT-3' |
| *KIF4A* | QT00037401 | | |
| *MIS18A* | QT00040502 | | |
| *SDHA* | | 5'-TGG GAA CAA GAG GGC ATC TG-3' | 5'-CCA CCA CTG CAT CAA ATT CAT G-3' |
| *RPS18* | | 5'- GAG GAT GAG GTG GAA CGT GT-3' | 5'-TCT TCA GTC GCT CCA GGT CT-3' |
| *TERT* | | 5'- GTC TGG AGC AAG TTG CAA AG-3' | 5'- CGA TGC TGC CTG ACC TCT-3' |
| *UHRF1* | QT00018256 | | |

[1] Qiagen, Hilden, Germany

### Supplementary Table S5. Antibodies used for western blotting.

| Target protein | Antibody source / Type | Supplier ID & Clone |
|---|---|---|
| Human **HDAC1** | Rabbit monoclonal | Abcam[1], clone 10E2, ab7028 |
| Human **HDAC2** | Mouse monoclonal | Santa Cruz Biotechnology[2], clone 3F3, sc-81599 |
| Human **GAPDH** | Mouse polyclonal | Merck Millipore[3], mouse monoclonal; clone 6C5, MAB374 |
| | Peroxidase-conjugated goat anti-mouse IgG | Dianova GmbH[4], 111-035-003 |
| | Peroxidase-conjugated goat anti-rabbit IgG | Dianova GmbH[4], 115-035-003 |

1 Cambridge, UK
2 Dallas, TX, U.S.A.
3 Darmstadt, Germany
4 Hamburg, Germany

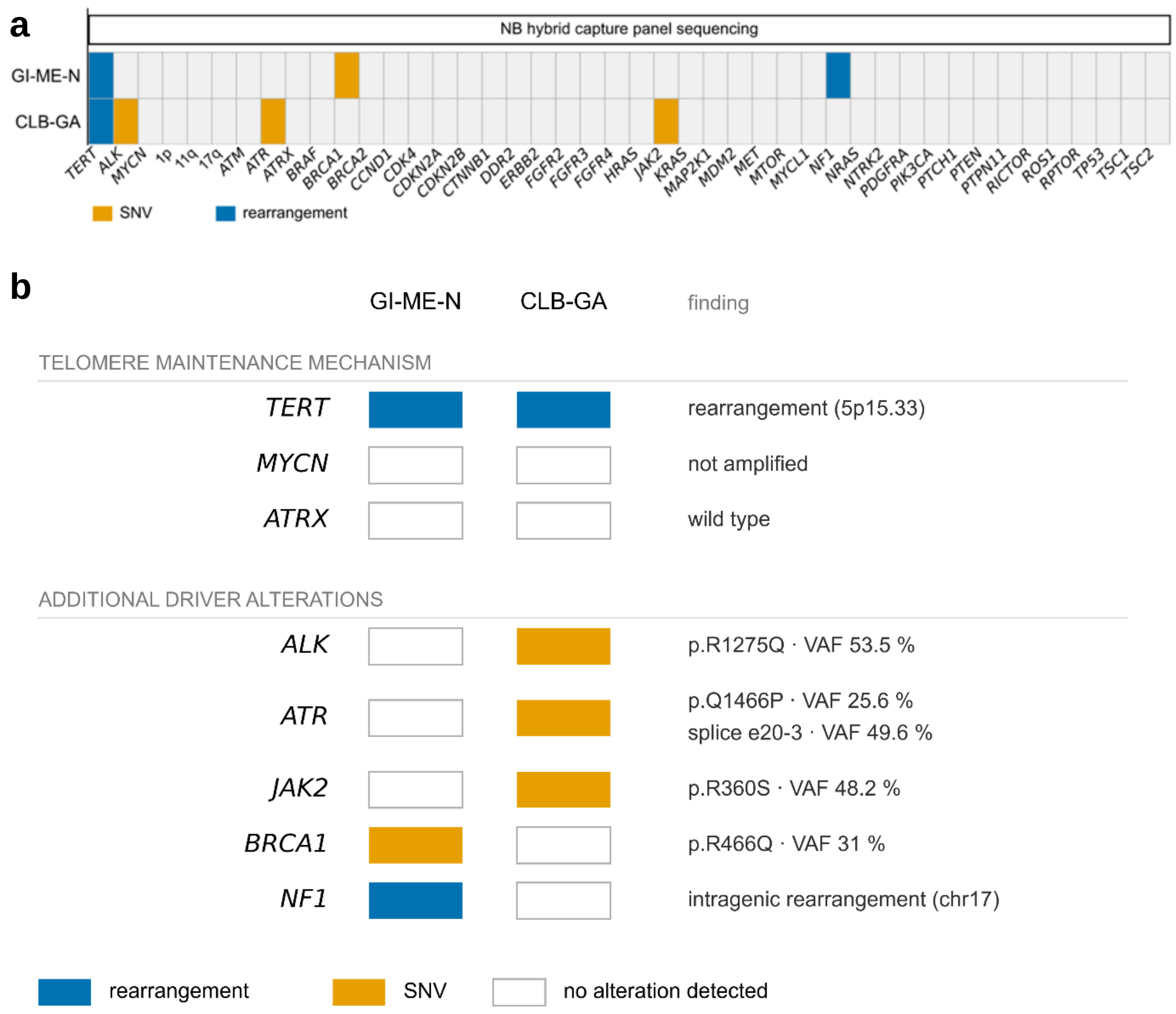


**Suppl. Fig. S1. Genomic panel profiling of *TERT*-rearranged GI-ME-N and CLB-GA neuroblastoma cell lines.** **a**, Heatmap displaying hybrid capture panel sequencing[22] results across key neuroblastoma cancer driver genes in GI-ME-N and CLB-GA cells. **b**, Detailed overview of the telomere maintenance mechanisms (*upper panel*) and variant allele frequencies (VAF) of additional driver alterations (*lower panel*) identified in the GI-ME-N and CLB-GA cell lines. The term "no alteration detected" refers to the specific genes and genomic regions indicated.

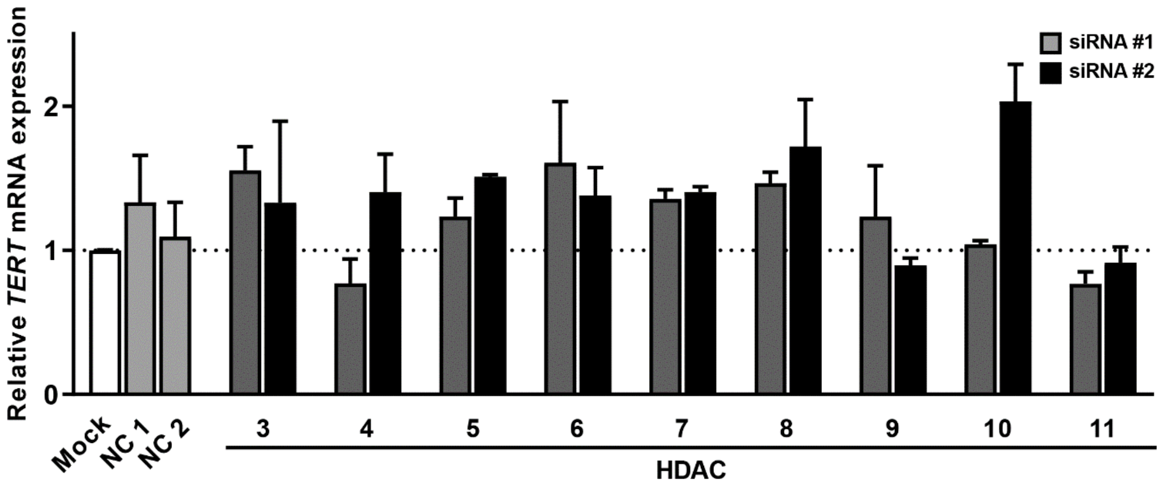


**Suppl. Fig. S2. Influence of HDAC family members 3 – 11 on *TERT* mRNA expression in GI-ME-N cells.** Shown are relative *TERT* mRNA levels measured by qRT-PCR at 120 h post-transfection with siRNAs directed individually against HDAC family members 3 - 11 or non-silencing negative controls. Data represent mean ± SD (n ≥ 3). Mock, transfection reagent only; NC, negative control siRNA.

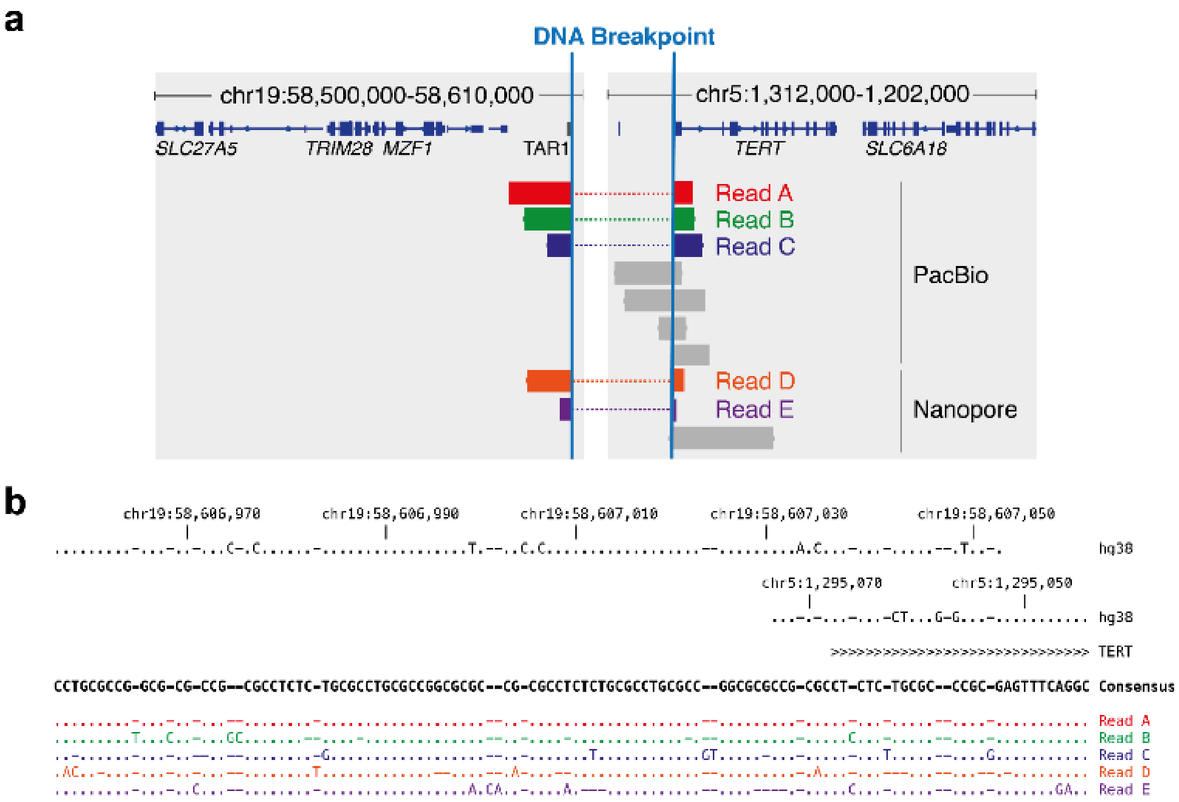


**Suppl. Fig. S3. Molecular *TERT* breakpoint characterization. a**, Schematic representation of the genomic breakpoint joining chr19 (SLC27A5/TAR1 region) and chr5 (*TERT* locus) derived from Pacific Biosciences (PacBio) and Oxford Nanopore long-read sequencing alignments in GI-ME-N cells. **b**, Nucleotide sequence alignment of consensus breakpoint reads against the human reference genome (hg38).

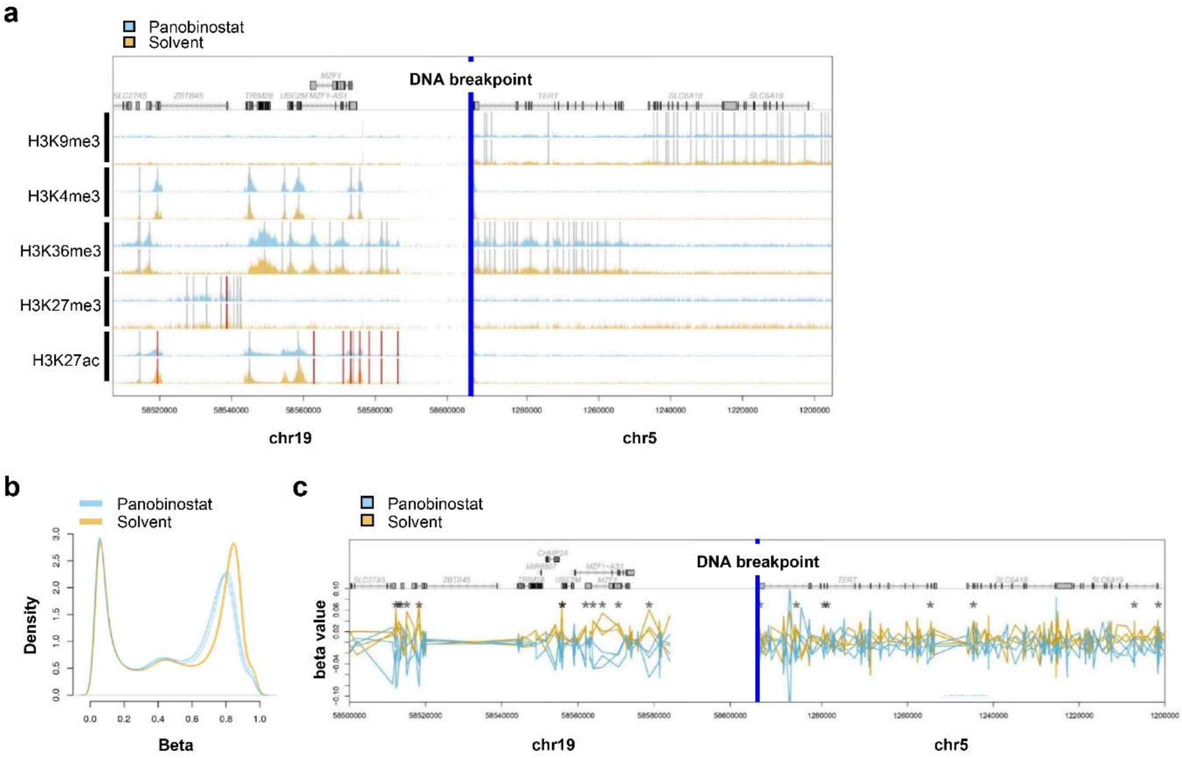


**Suppl. Fig. S4. Epigenetic histone modifications and DNA methylation across the *TERT* translocation breakpoint upon panobinostat treatment. a**, ChIP-seq signal tracks for repressive (H3K9me3, H3K27me3) and active (H3K4me3, H3K36me3, H3K27ac) histone marks across the chr19-chr5 translocation breakpoint in GI-ME-N cells treated for 18 h with 20 nM Panobinostat or solvent control. **b**, Density plot of genome-wide DNA methylation beta values in panobinostat- versus solvent-treated GI-ME-N cells measured by Illumina Infinium Human MethylationEPIC BeadChip. **c**, Locus-specific DNA methylation beta values across chr19 and chr5 breakpoint regions showing an unaltered local CpG methylation status. * $P < 0.05$.

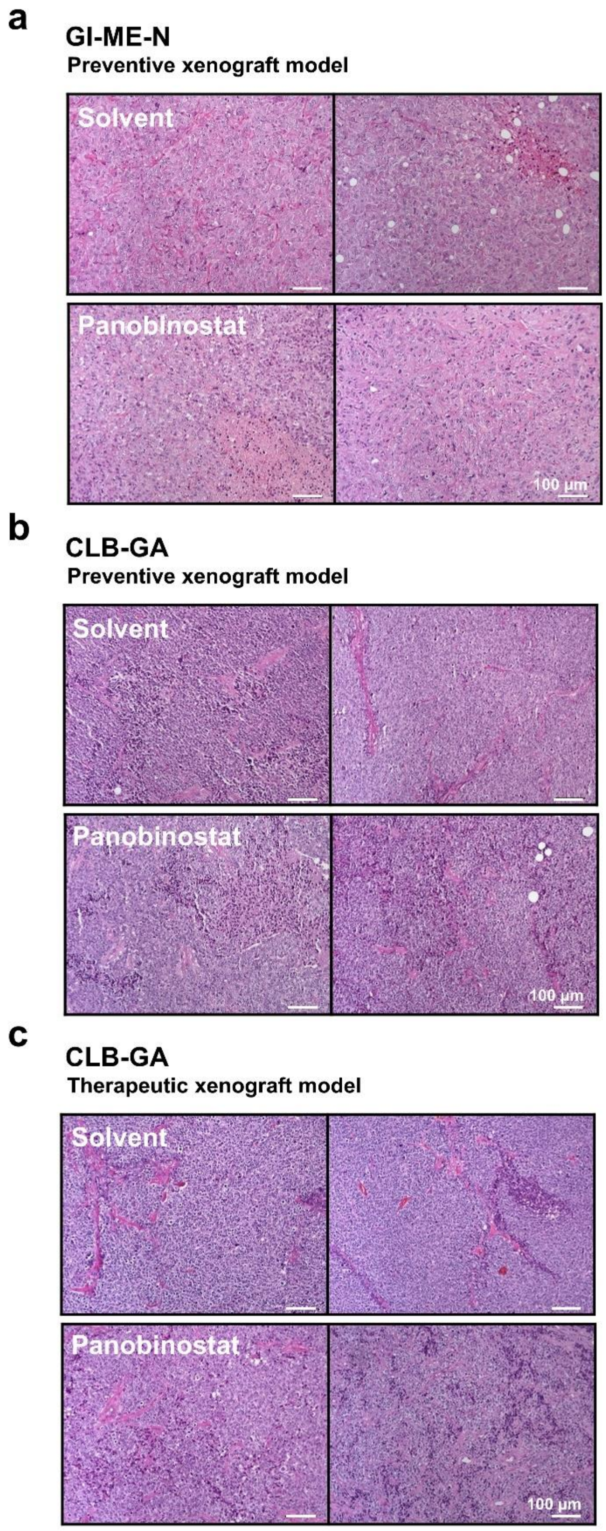


**Supplementary Fig. S5**. **Histopathological evaluation of xenograft tumor tissue following panobinostat treatment.** Representative hematoxylin and eosin (H&E) stained histology sections of harvested xenograft tumor tissues from GI-ME-N preventive (**a**), CLB-GA preventive (**b**) and CLB-GA therapeutic models (**c**), demonstrating the high morphological tumor cell purity (>90% tumor cell content) typical of patient-derived neuroblastoma xenografts in mice across treatment and control arms. Scale bar, 100 µm.

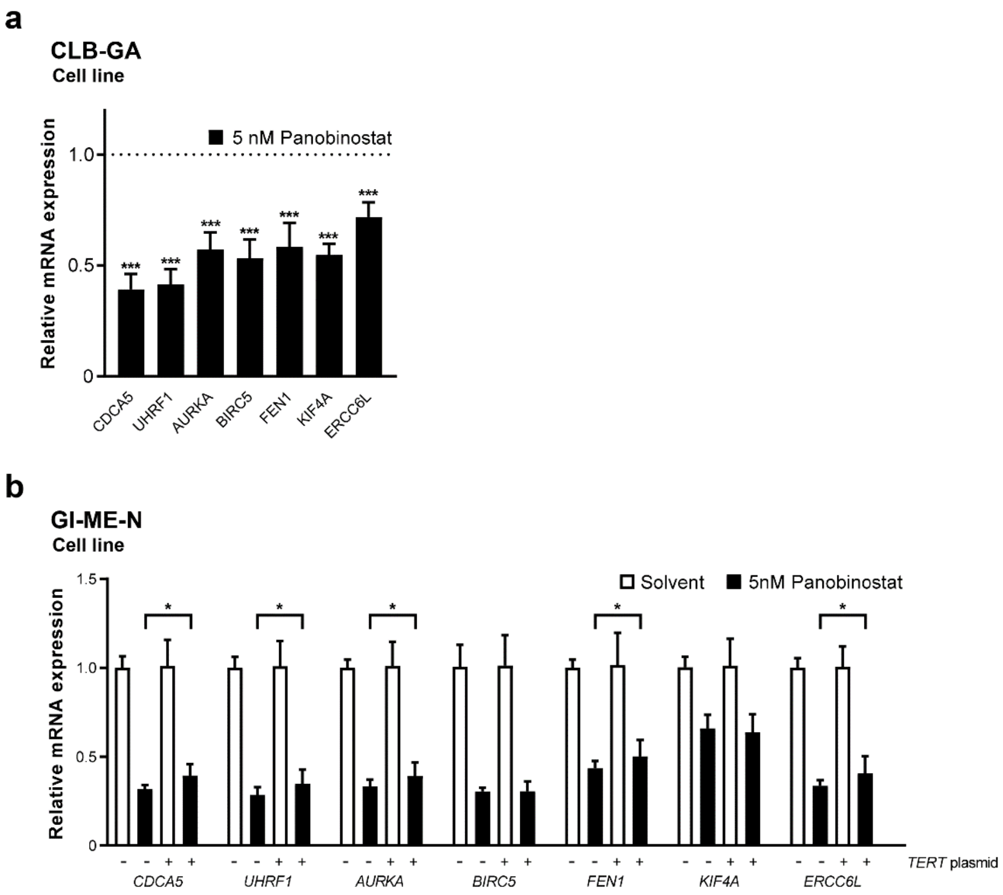


**Suppl. Fig. S6. Quantitative RT-PCR validation of core *TERT*-associated signature gene downregulation and genetic rescue upon panobinostat treatment. a**, Relative mRNA transcript levels of representative core 50-gene TERT-associated signature targets measured by qRT-PCR following panobinostat treatment (5 nM) versus solvent control in *TERT*-rearranged CLB-GA neuroblastoma cells, confirming the downregulation observed by RNA-Seq analysis. **b**, Relative mRNA transcript levels of representative signature targets measured by qRT-PCR in cells transiently transfected with ectopic *TERT* cDNA plasmid (+) or empty control vector (-) upon panobinostat treatment, demonstrating genetic rescue of signature gene expression following *TERT* re-expression. Data represent mean ± SD (n = 3). **P* < 0.05; ****P* < 0.001.

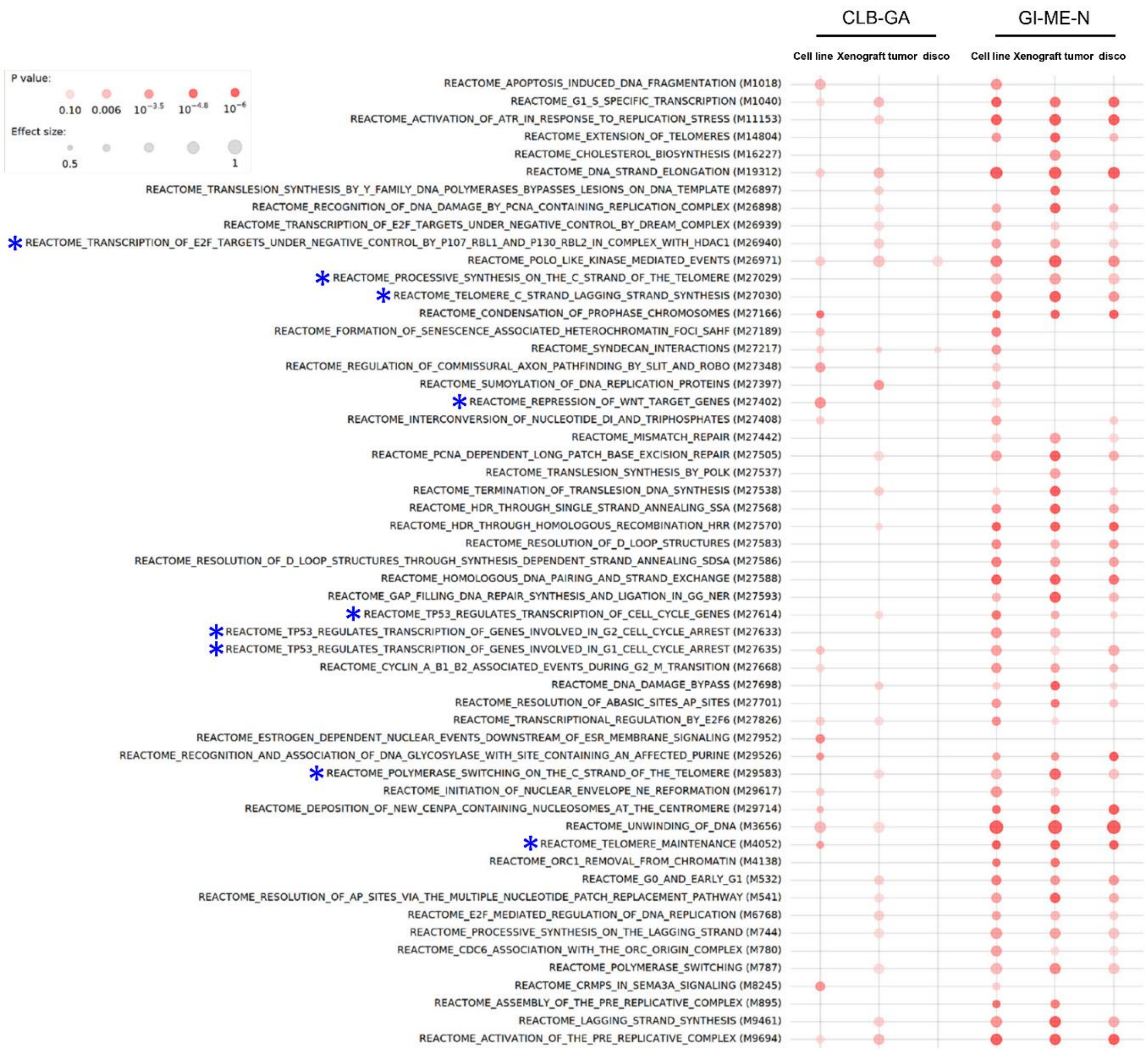


**Suppl. Fig. S7. Transcriptomic profiling and REACTOME pathway enrichment analysis in CLB-GA and GI-ME-N cell lines, preventive xenograft tumors and the discovery cohort.** RNA sequencing (RNA-seq) data and REACTOME pathway enrichment analysis comparing panobinostat-treated versus solvent-controlled samples across CLB-GA and GI-ME-N neuroblastoma models, encompassing cell lines, xenograft tumors from the preventive mouse model and the discovery cohort (representing the combined cell line and tumor dataset). Blue asterisks indicate pathways and gene sets directly linked to telomere maintenance and function.
